# Changing protein-DNA interactions promote ORC binding site exchange during replication origin licensing

**DOI:** 10.1101/2023.06.16.545300

**Authors:** Annie Zhang, Larry J. Friedman, Jeff Gelles, Stephen P Bell

**Author notes:** Co-corresponding authors: Stephen P. Bell, Jeff Gelles.

## Abstract

During origin licensing, the eukaryotic replicative helicase Mcm2-7 forms head-to-head double hexamers to prime origins for bidirectional replication. Recent single-molecule and structural studies revealed that one molecule of the helicase loader ORC can sequentially load two Mcm2-7 hexamers to ensure proper head-to-head helicase alignment. To perform this task, ORC must release from its initial high-affinity DNA binding site and “flip” to bind a weaker, inverted DNA site. However, the mechanism of this binding-site switch remains unclear. In this study, we used single-molecule Förster resonance energy transfer (sm-FRET) to study the changing interactions between DNA and ORC or Mcm2-7. We found that the loss of DNA bending that occurs during DNA deposition into the Mcm2-7 central channel increases the rate of ORC dissociation from DNA. Further studies revealed temporally-controlled DNA sliding of helicase-loading intermediates, and that the first sliding complex includes ORC, Mcm2-7, and Cdt1. We demonstrate that sequential events of DNA unbending, Cdc6 release, and sliding lead to a stepwise decrease in ORC stability on DNA, facilitating ORC dissociation from its strong binding site during site switching. In addition, the controlled sliding we observed provides insight into how ORC accesses secondary DNA binding sites at different locations relative to the initial binding site. Our study highlights the importance of dynamic protein-DNA interactions in the loading of two oppositely-oriented Mcm2-7 helicases to ensure bidirectional DNA replication.

**Significance Statement:** Bidirectional DNA replication, in which two replication forks travel in opposite directions from each origin of replication, is required for complete genome duplication. To prepare for this event, two copies of the Mcm2-7 replicative helicase are loaded at each origin in opposite orientations. Using single-molecule assays, we studied the sequence of changing protein-DNA interactions involved in this process. These stepwise changes gradually reduce the DNA-binding strength of ORC, the primary DNA binding protein involved in this event. This reduced affinity promotes ORC dissociation and rebinding in the opposite orientation on the DNA, facilitating the sequential assembly of two Mcm2-7 molecules in opposite orientations. Our findings identify a coordinated series of events that drive proper DNA replication initiation.

## Introduction

Eukaryotic chromosomal replication initiates by building two oppositely-oriented replication forks at origins of replication. During G1, the core component of the replicative helicase, the Mcm2-7 complex, is assembled around origins in a process termed origin licensing or helicase loading. This process marks all potential origins of replication and a subset of the loaded helicases are activated in S phase to form the core of the replication machinery (1). To ensure bidirectional replication, each pair of Mcm2-7 helicases must be assembled around origin DNA in a head-to-head orientation (2–6).

In budding yeast, a “one-loader” helicase loading mechanism has been proposed in which one molecule of the helicase-loader, the origin recognition complex (ORC), coordinates the loading of both Mcm2-7 hexamers (7–9). In this model, ORC begins helicase loading by binding origin DNA and recruiting Cdc6 (Fig. 1A, step 1) (10, 11). The ORC-Cdc6-DNA complex recruits a Mcm2-7-Cdt1 complex (Fig. 1A, step 2) to form the transient <u>O</u>RC-<u>C</u>dc6-<u>C</u>dt1-<u>M</u>cm2-7 (OCCM) intermediate (Fig. 1A, step 3). Cdc6 and then Cdt1 are subsequently released in a strict sequential order (Fig. 1A, steps 4-5) (7). Upon Cdt1 dissociation, the initial ORC-Mcm2-7 interaction is broken, allowing this ORC molecule to “flip” to the opposite side of the helicase and form a distinct <u>M</u>cm2-7-<u>O</u>RC (MO) intermediate (Fig. 1A, step 6) (8, 9). MO formation mediates stable closing of the Mcm2/5 gate, a gap between subunits Mcm2 and Mcm5 that allows access of DNA to the Mcm2-7 central channel, such that the helicase stably encircles DNA (8, 12). The resulting MO complex is required to recruit and load a second helicase in the opposite orientation (8, 9). Once recruited, the second helicase rapidly forms the head-to-head interactions with the first helicase.

**Figure 1:**
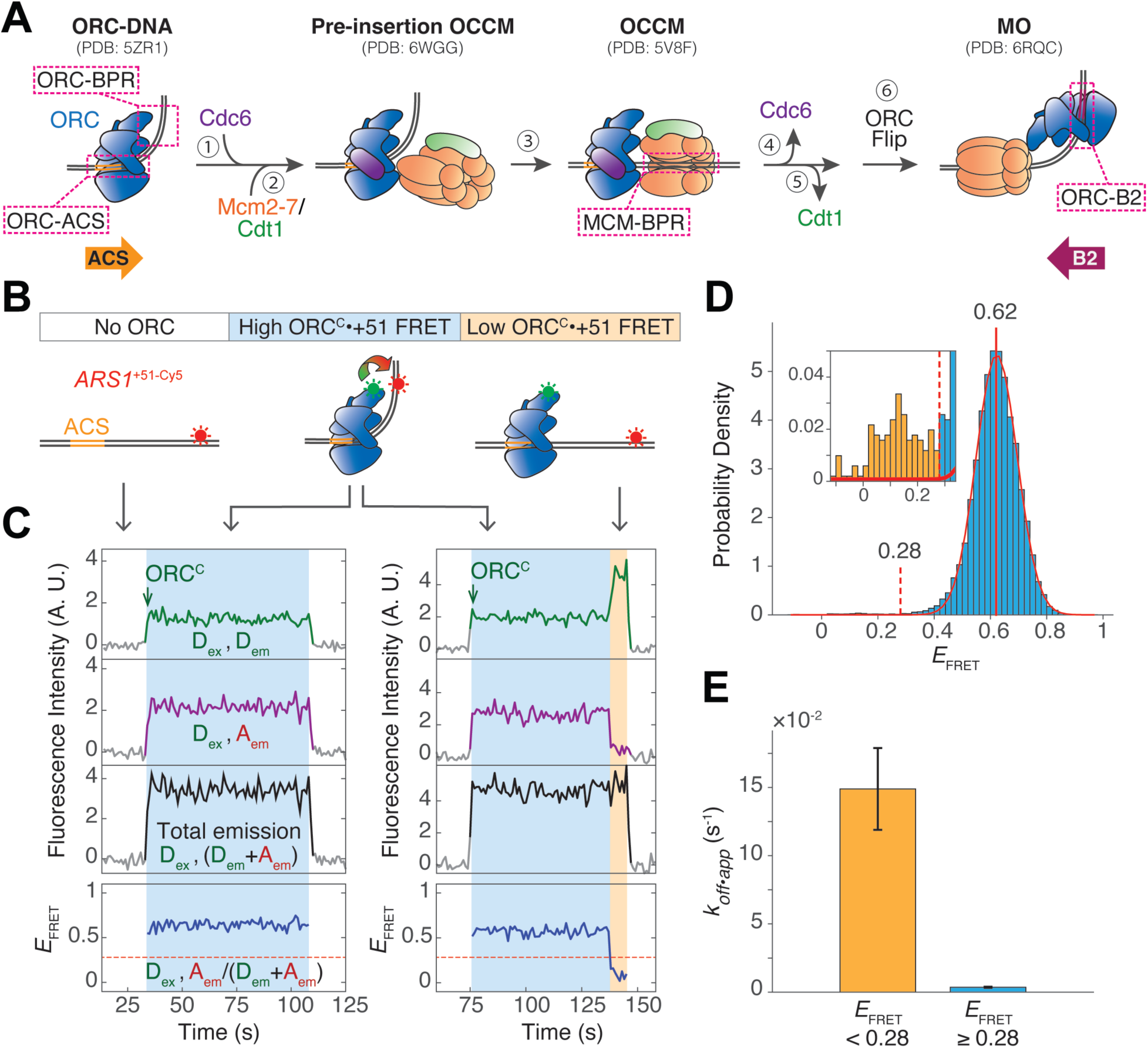
ORC introduces a stable bend in DNA that increases its stability on DNA. **A.** Model of helicase-loading from initial ORC recruitment onto origin DNA to MO formation. Illustrations based on cryo-EM structures of helicase-loading intermediates are labeled with PDB identifiers. Pink dashed boxes: protein-DNA interactions. Colored arrows on the bottom show the relative orientation of the ACS and B2 elements. Further details are given in the text. **B.** Schematic of ORC^C^•+51 sm-FRET experiment that monitors ORC-BPR interaction. ORC^C^•+51 FRET (curved arrow) is high when the complex is in the bent state, and low in the unbent state. **C.** ORC^C^•+51 FRET assay records at two individual DNA molecules. Top plot: donor emission during donor excitation (green, D_ex_, D_em_). Green arrow indicates ORC^C^ arrival on DNA. Second plot: acceptor emission during donor excitation (purple, D_ex_, A_em_). Third plot: total emission during donor excitation (black, D_ex_, (D_em_+A_em_)). Bottom plot: effective FRET efficiency (*E*_FRET_) (blue, D_ex_, A_em_ /(D_em_+A_em_)). Grey segments represent background signal when no fluorescent ORC is colocalized with DNA; segments of the intensity line plot during ORC colocalization are colored. Colored background indicates ORC-DNA in the high *E*_FRET_ bent state (blue, as in B), or in the low *E*_FRET_ unbent state (orange, as in B). A red dashed line at *E*_FRET_ = 0.28 separates the high and low *E*_FRET_ states. Additional traces are shown in *SI Appendix*, Fig S1B. **D.** Histogram of *E*_FRET_ values during each frame when ORC was present on a DNA. Center *E*_FRET_ value from a one-component Gaussian fit (red curve; *SI Appendix*, Table S1) is 0.62 ± 0.001 (red vertical line). Inset: magnified histogram. *E*_FRET_ values above (blue) or below (yellow) the 0.28 threshold (red dashed line) were shown. Data are from 25,388 acquired frames during 186 ORC colocalization events. **E.** Apparent dissociation rate of ORC from DNA (*k_off•app_*) when ORC is in the low *E*_FRET_ state (< 0.28) or the high *E*_FRET_ state (≥ 0.28). Error bars indicate the standard error. Number of video frames in low *E*_FRET_ state = 141; number of frames in high *E*_FRET_ state = 25,247.

In addition to protein-protein interactions, ORC forms extensive but changing interactions with DNA (Fig. 1A, dashed boxes). In budding yeast, the primary ORC binding site is defined by the ARS consensus sequence (ACS) and B1 element (11, 13). ORC encircles the ACS as an open ring, forming additional interactions with DNA including and beyond the B1 element to create a strong ∼80° bend in DNA (14). Structural studies have captured a “pre-insertion OCCM” intermediate with bent DNA that contains all helicase-loading proteins (Fig. 1A) (15). However, the bent DNA straightens and is inserted in the Mcm2-7 open ring in the later OCCM intermediate (16). These intermediates together suggest that a segment of DNA flanking the bend, which we refer to as the bend-proximal region (BPR) (Fig. 1A), is transferred from ORC to Mcm2-7, allowing the DNA bend to straighten as the BPR enters the Mcm2-7 central channel. Although molecular dynamics models have simulated the DNA insertion process (15), DNA unbending and deposition into Mcm2-7 have yet to be monitored in real time. In addition, the trigger(s) for this change in ORC-DNA interactions has not been defined.

For one ORC to load both Mcm2-7 helicases at an origin, ORC has to switch between two oppositely-oriented DNA binding sites. Natural yeast origins include at least one B2 element, each of which includes a partial-match to the ACS but in the inverted orientation (Fig. 1A, colored arrows) (17–20). As a result, the B2 element is proposed to serve as a weaker secondary binding site for ORC. Studies with artificial origins showed that a second inverted binding site is essential for robust helicase loading (21). Interestingly, B2 elements are found at variable distances from the ACS (17, 19, 22). A key question raised by the “one-loader” hypothesis for helicase loading is how ORC releases from its strong initial binding site to bind a weaker, oppositely-oriented secondary binding site a variable distance away.

Combining colocalization single-molecule spectroscopy (CoSMoS) (23, 24) and single-molecule Förster resonance energy transfer (sm-FRET) (25), we monitored a variety of interactions between ORC and Mcm2-7 with the origin DNA and identified key triggers for changes in the observed protein-DNA interactions. We demonstrated that the ORC-DNA complex predominantly contains bent DNA, but infrequently transitions to a less stable conformation with straightened DNA. Cdc6 binding does not change the bent state of the ORC-bound DNA but does decrease the dissociation of ORC from DNA. Although the OCCM is static on the DNA, upon loss of Cdc6 (Fig. 1A, step 4) we observe changes in sm-FRET supporting sliding of the <u>O</u>RC-<u>C</u>dt1-<u>M</u>cm2-7 (OC_1_M) complex. Based on our data, we propose that the sequential events of DNA unbending, Cdc6 release, and DNA sliding progressively reduce ORC stability on DNA to facilitate ORC dissociation from DNA and flipping to bind the inverted B2 element.

## Results

### The ORC-DNA complex predominantly exists in a bent DNA state

To understand how and when origin DNA changes its conformation during helicase loading, we developed a single-molecule FRET (sm-FRET) assay to monitor the dynamics of ORC-induced DNA bending. Previous cryo-EM structures showed that the C-terminal face of ORC, particularly the C-terminal domain of Orc6, is in close proximity to the bend-proximal region (BPR) of origin DNA (14). To position a donor fluorophore close to the BPR, we generated an ORC fluorescently labeled on its C-terminal face (ORC^C^). To this end, we deleted residues 1-266 of Orc6 and attached a fluorophore to the truncated N-terminus. Although defective in recruiting a second Mcm2-7 (26), truncation of this region in Orc6 does not compromise OCCM formation in ensemble helicase-loading assays (*SI Appendix*, Fig. S1A). An acceptor fluorophore was coupled to DNA at an internal position 51 bp from the *ARS1* origin ACS (*ARS1*^+51-Cy5^). This position was chosen to maximize proximity to the Orc6 label when ORC is bound to the BPR and minimize their proximity when the DNA is straight (Fig. 1B). We refer to the FRET between this donor-acceptor pair as “ORC^C^•+51” FRET (Fig. 1B), and the corresponding interaction between ORC and the BPR as the “ORC-BPR” interaction (Fig. 1A, first panel, dashed box). Using total internal reflection fluorescence (TIRF) microscopy, we monitored the colocalization of ORC^C^ in solution with surface-tethered origin DNA (23). Throughout each ORC-DNA colocalization event, we measured effective ORC^C^•+51 FRET efficiency (*E*_FRET_) to examine ORC-induced DNA bending.

When ORC arrived on DNA, we consistently observed a high ORC^C^•+51 *E*_FRET_ state, indicating ORC was bound to the ACS and BPR and that the bound DNA was bent (blue background in Fig. 1C-D). The majority of ORC^C^•+51 *E*_FRET_ values were centered at 0.62 (Fig. 1D). Although they represented less than 0.5% of the time of ORC-DNA colocalization, we also observed short periods of lower *E*_FRET_ values (< 0.28, orange background in Fig. 1C-D, *SI Appendix*, Fig. S1B). These low *E*_FRET_ values were not caused by photobleaching, as we examined the DNA-coupled fluorophore by acceptor excitation after the experiment and restricted analysis to DNA molecules that maintained fluorescence to the end of the experiment. In addition, control experiments showed that these low *E*_FRET_ values were not caused by ORC sliding along the DNA to move away from the ACS (*SI Appendix*, Fig. S2). We excluded a small fraction (approximately 2%) of ORC molecules that did not show a high *E*_FRET_ signal at any time during the DNA colocalization, as these likely represent nonspecific ORC binding. The poor fit of a single Gaussian model at low *E*_FRET_ values (Fig. 1D, red curve in inset) suggests the presence of distinct populations at the bent or unbent conformational states. We conclude that the low *E*_FRET_ values reflect an unbent state in which the ORC remains bound at the ACS but the ORC-BPR interaction is lost (Fig. 1B). Addition of Cdc6 increases ORC stability on DNA, leading to a near two-fold decrease in its apparent dissociation rate from 0.0131 s^-1^ to 0.0073 s^-1^ (*SI Appendix*, Fig. S1C, Table S1). However, Cdc6 did not shift the ORC^C^•+51 *E*_FRET_ distribution between the bent and unbent states, consistent with Cdc6 not significantly altering the ORC-BPR interaction (*SI Appendix*, Fig. S1C) (15, 27). Thus, ORC-bound DNA is predominantly in the bent state with infrequent transitions to the unbent state.

We frequently observed the low *E*_FRET_ unbent state immediately prior to ORC dissociation from DNA (Fig. 1C, *SI Appendix*, Fig. S1B). Thus, we asked whether DNA unbending increases the rate of ORC dissociation from DNA. Indeed, ORC in the low *E*_FRET_ state (< 0.28) dissociates more than 40 times faster from DNA than ORC in the high *E*_FRET_ state (≥ 0.28) (Fig. 1E). These data indicate that the ORC-BPR interaction increases ORC stability on DNA.

As a further test of the ORC-BPR contribution to the kinetic stability of the ORC-DNA complex, we mutated 9 amino acids in the ORC region that interacts with the phosphate backbone of the BPR (14) (*SI Appendix*, Fig. S1D). When these mutations are incorporated into labeled ORC (ORC^C-BPR^), we observed a limited decrease in *E*_FRET_ (centered at 0.52, *SI Appendix*, Fig. S1E), suggesting that these interactions contribute to but are not sufficient for ORC-induced DNA bending. Nevertheless, ORC^C-BPR^ stability on DNA is notably reduced, reflected by a significant increase in its apparent dissociation rate (0.0849 s^-1^) compared to WT (0.0131 s^-1^, *SI Appendix*, Table S1). We conclude that the kinetic stability of ORC on DNA is strongly dependent on the ORC-BPR interaction, as a partial loss of this interaction results in a strong reduction in ORC-DNA stability.

### Mcm4 and Mcm6 winged-helix domains trigger DNA unbending

After Mcm2-7 recruitment, the BPR disengages from ORC and is inserted into the helicase central channel (16). To understand the transition of the BPR DNA from being bound to ORC-Cdc6 to being deposited into the helicase, we investigated the kinetics of DNA unbending during helicase loading. We included Cdc6 and labeled Mcm2-7^4N-650^/Cdt1 in our experiment to simultaneously monitor ORC-BPR interactions and Mcm2-7 recruitment. To avoid using red-excited fluorophores on both DNA and Mcm2-7, we replaced the DNA-coupled acceptor with the fluorescence quencher Black Hole Quencher-2 (BHQ-2) at the same DNA site (*ARS1*^+51-BHQ2^). In this experiment, DNA bending induced by ORC-BPR interaction results in quenching of green-excited ORC^C^ fluorescence, and DNA unbending due to loss of the ORC-BPR interaction leads to unquenching (Fig. 2A). Using increase in green-excited fluorescence as a marker for DNA unbending, we consistently detected a temporal delay between Mcm2-7 arrival (Fig. 1A, step 2) and DNA unbending (Fig. 2B, *SI Appendix*, Fig. S3). Unbending occurs 3.9 ± 0.3 s (mean ± SEM) after Mcm2-7 arrival, which is significantly earlier (Fig. 2C) than Cdc6 release (Fig. 1A, step 4) (7). Given the temporal delay between Mcm2-7 recruitment and DNA unbending, we explored the possibility of additional regulatory steps between these two events.

**Figure 2:**
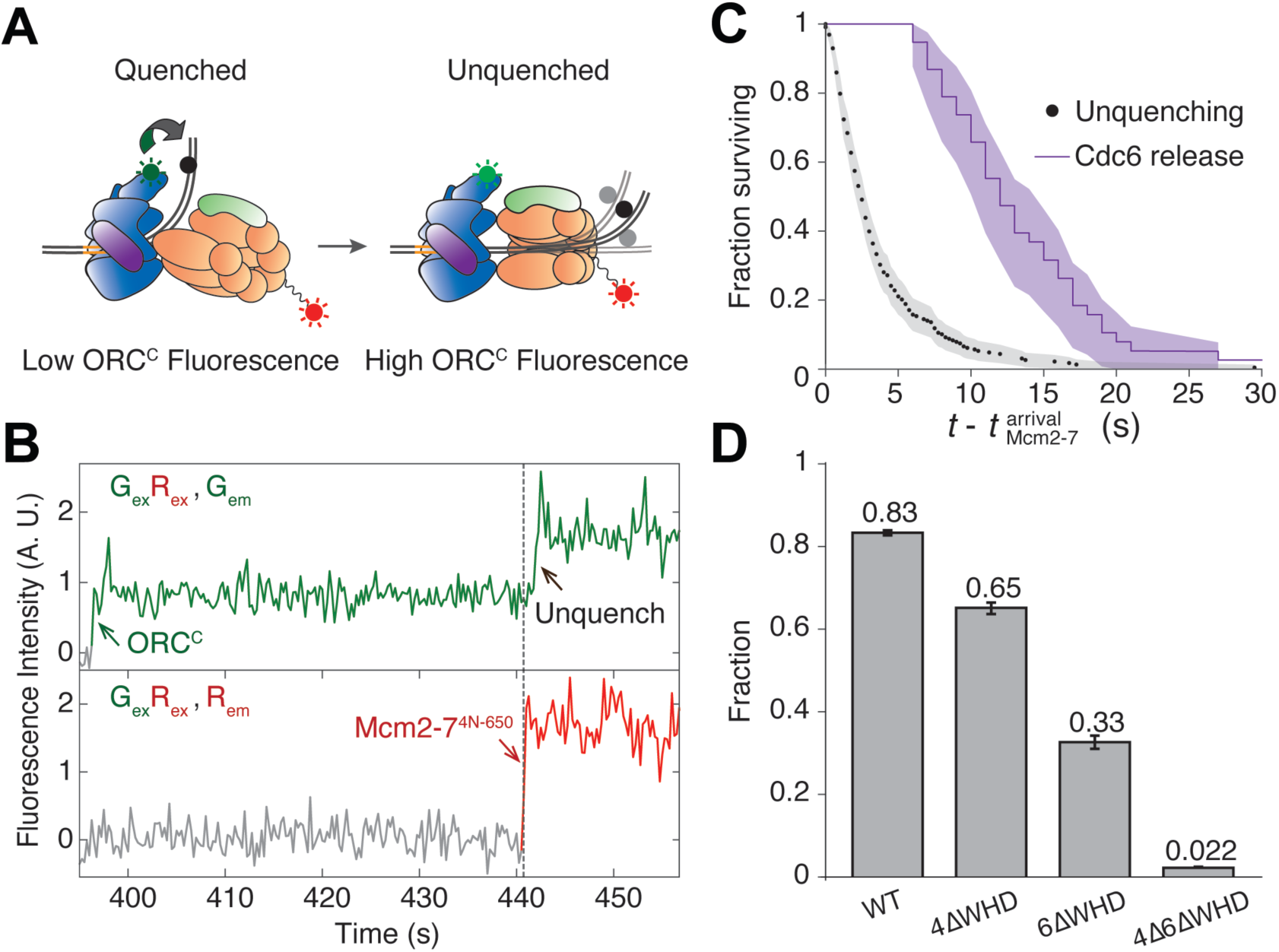
DNA unbending occurs rapidly after Mcm2-7 recruitment and is driven by winged-helix domains of Mcm4 and Mcm6. **A.** Schematic of fluorescence quenching experiment monitoring ORC-BPR interaction during the loading of the first Mcm2-7 helicase. Mcm2-7 is labeled with a red-excited fluorophore at the unstructured Mcm4 N-terminal tail (Mcm2-7^4N-650^). The black dot represents the quencher placed at the +51 position on *ARS1*. **B.** Representative record from the experiment in (a). Top plot: green emission from green and red simultaneous excitation (G_ex_R_ex_, G_em_). Green arrow marks ORC^C^ association with DNA, and black arrow marks unquenching of ORC-associated fluorophore. Bottom plot: red emission from green and red simultaneous excitation (G_ex_R_ex_, R_em_). Dashed line and red arrow marks Mcm2-7^4N-650^ arrival. Time resolution = 0.25 s. Additional examples are shown in *SI Appendix*, Fig. S3. **C.** Survival functions comparing the interval between Mcm2-7 arrival and unquenching (i.e. DNA unbending, black) and the interval between Mcm2-7 arrival and Cdc6 release (purple). Shading: 95% confidence interval (CI). 228 DNA unbending and 82 Cdc6 release events are plotted. The Cdc6 data was originally reported in Ticau et al. 2015, Fig 4D. **D.** Fraction of Mcm2-7 colocalization events during which DNA unbent. Error bars: standard error.

Previous biochemical and structural studies suggest that Mcm2-7 association with the ORC-Cdc6-DNA complex is a multistep process. After ORC-Cdc6 initially recruits Mcm2-7 via the winged-helix domains (WHDs) of Mcm3 and Mcm7, additional interactions are formed between ORC and Mcm2-7 (15, 28, 29). In particular, the C-terminal WHDs of Mcm4 and Mcm6 establish additional interactions with ORC between initial helicase recruitment and OCCM formation. To address if these new interactions promote DNA unbending, we created red-excited Mcm2-7 mutants labeled at Mcm4 N-terminus that lack the WHDs in Mcm4 (4ΔWHD), Mcm6 (6ΔWHD), or both subunits (4Δ6ΔWHD). We tested each of these mutants in the ORC^C^•+51 unquenching assay. Using unquenching as a marker for DNA unbending, we determined the fraction of Mcm2-7-DNA colocalization events that resulted in DNA unbending. Deletion of either the Mcm4 or Mcm6 WHDs showed reduced fractions of Mcm2-7 colocalization events that displayed unbending, with the Mcm6 WHD deletion having the stronger effect (Fig. 2D). Simultaneous deletion of the Mcm4 and Mcm6 WHDs almost completely abolished DNA unbending (Fig. 2D). Although these mutations could delay DNA unbending or lead to Mcm2-7 dissociation before DNA unbending occurs, the time from Mcm2-7 arrival to DNA unbending was not changed by the mutations and this time is significantly faster than Mcm2-7 dissociation for each mutant (*SI Appendix*, Fig. S4A). Thus, these findings strongly suggest that interactions made by ORC with the Mcm4 and Mcm6 WHDs trigger DNA unbending, coordinating this event with proper ORC-Mcm2-7 positioning.

### DNA is deposited rapidly into the central channel of Mcm2-7 after DNA unbending

Structural studies of the pre-insertion OCCM intermediate show that ORC holds the BPR directly above the Mcm2/5 gate to position BPR for entry into the Mcm2-7 ring (Fig. 1A) (15). This finding suggests that DNA unbending would lead to rapid DNA deposition into the Mcm2-7 central channel. To monitor DNA deposition and test this hypothesis, we developed a FRET-based assay to detect MCM-BPR interactions (Fig. 1A) using an Mcm2-7-coupled donor at the N-terminus of Mcm3 (Mcm2-7^3N-550^) and the same DNA-coupled acceptor at the +51 position within BPR (*ARS1*^+51-Cy5^). Instead of ORC^C^, unlabeled ORC with full-length Orc6 was used, allowing for complete helicase loading. In this assay, a high “MCM•+51” *E*_FRET_ is expected following DNA insertion into the central channel of the first Mcm2-7 (Fig. 3A). Consistent with this prediction, we observed periods of high *E*_FRET_ shortly after first Mcm2-7 arrival (Fig. 3B). In 135 Mcm2-7 colocalization events with DNA, most of the complexes initially had ORC-engaged BPR, as evidenced by a low MCM•+51 *E*_FRET_ distribution (centered at 0.12) 1 - 2 s after Mcm2-7 association (Fig. 3C). In contrast, 10 - 11 s after Mcm2-7 arrival, the MCM•+51 *E*_FRET_ values were higher (centered at 0.55), implying that DNA had been deposited into the Mcm2-7 channel (Fig. 3C). Using the increase to high *E*_FRET_ (> 0.35, see Methods) as a marker for DNA deposition, the average time between Mcm2-7 arrival and DNA deposition was 4.9 ± 0.7 s (Fig. 3D). Consistent with DNA deposition being dependent on DNA unbending, the Mcm2-7 WHD mutants that exhibited reduced DNA unbending (Fig. 2D) also exhibited reduced DNA deposition (*SI Appendix*, Fig. S4B). Similar distributions are observed for both time to unbending and time to deposition after Mcm2-7 arrival (Fig. 3D), suggesting a rapid transfer of the BPR DNA from ORC to the Mcm2-7 central channel.

**Figure 3:**
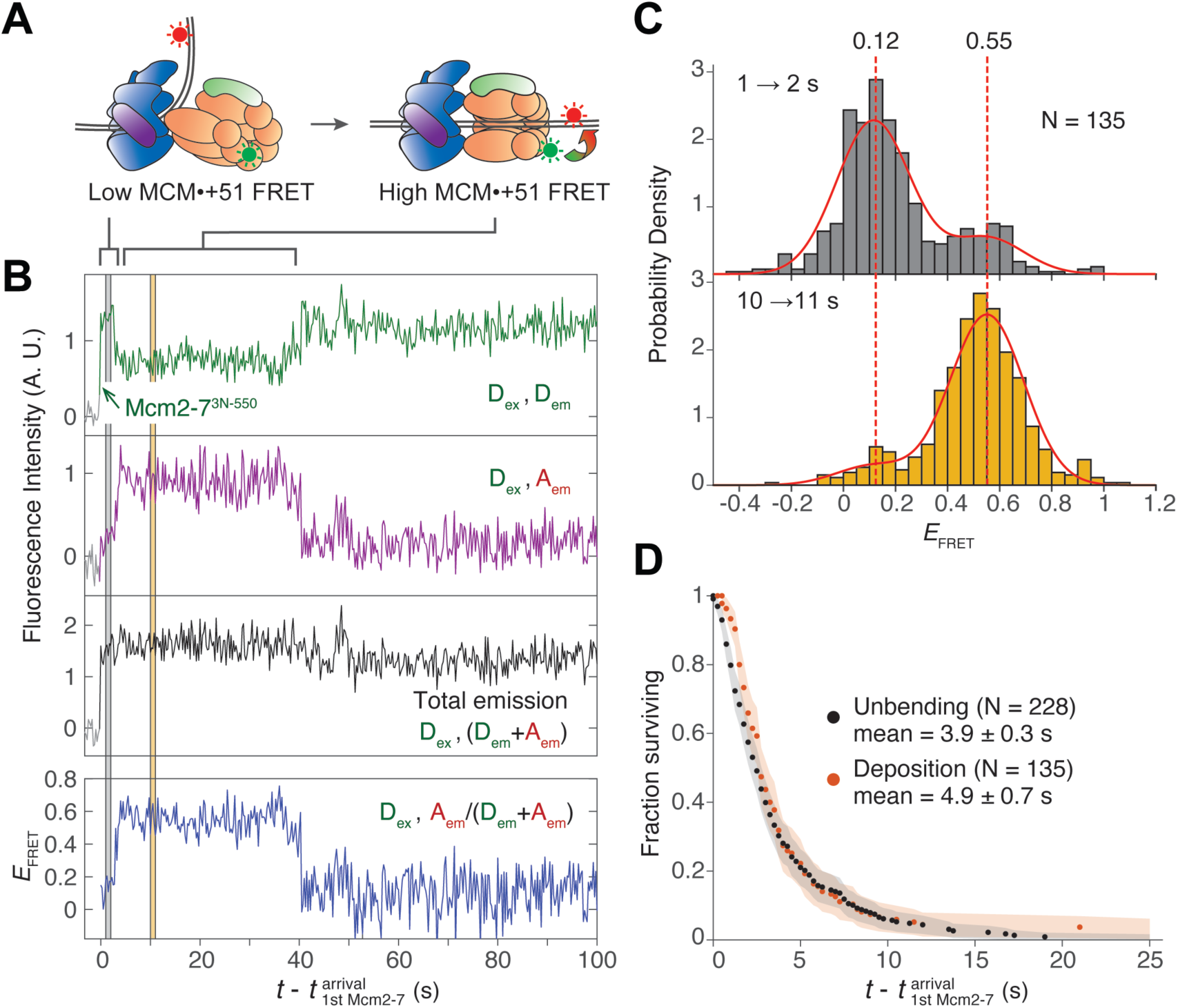
DNA deposition into the Mcm2-7 central channel exhibits a similar temporal distribution to DNA unbending. **A.** Schematic of the experiment monitoring DNA deposition into the Mcm2-7 central channel, which is indicated by high MCM•+51 *E*_FRET_. The FRET probes were Mcm2-7^3N-550^ and *ARS1*^+51-Cy5^. **B.** Example record from the experiment described in (a). Figure descriptions are as Fig. 1C, except that green arrow marks arrival of Mcm2-7 on DNA. Grey shading over the plots: 1s - 2s after Mcm2-7 arrival; yellow shading: 10 s - 11 s after Mcm2-7 arrival. Time resolution = 0.25 s. **C.** MCM•+51 *E*_FRET_ distribution during 1 – 2 s (grey, top plot) and 10 – 11 s after first Mcm2-7 arrival (yellow, bottom plot). Data were fitted by a global two-component Gaussian mixture model (solid red lines, *SI Appendix*, Table S1) with global component centers at 0.12 and 0.55 (red dashed lines). The time intervals correspond to the grey and yellow shadings in (B). N: number of Mcm2-7 colocalization events. **D.** Cumulative distribution of time intervals separating the first Mcm2-7 arrival and the subsequent DNA unbending (black, same as Fig. 2C) or DNA deposition (red). The mean time to reach unbending or deposition ± standard error of the mean are shown. Shaded areas: 95% CI; N: number of DNA-bound ORC or Mcm2-7 molecules that exhibited intensity or *E*_FRET_ change.

Shortly after the transition to high *E*_FRET_ upon Mcm2-7 arrival, we usually observed a decrease of *E*_FRET_ (e.g., Fig. 3B at 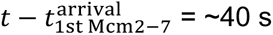). To ensure that this *E*_FRET_ decrease is associated with successful helicase loading, we limited our analysis to Mcm2-7-DNA colocalization events that resulted in high-salt-resistant, loaded Mcm2-7 hexamers (7, 30).

Using data from the experimental setup described in Fig. 3A, we plotted a two-dimensional heat map for these productive loading events to visualize MCM•+51 *E*_FRET_ values 0 - 100 s after the first Mcm2-7 arrival (Fig. 4A). We divided the heat map into four time windows (TW, Fig. 4A) and fit a two-component Gaussian model to the *E*_FRET_ distribution in each TW (Fig. 4B, red curves, *SI Appendix*, Table S1). The centers of the low and high Gaussian components in TW1 and TW2 were not significantly different (Fig. 4B, *SI Appendix*, Table S1). Consistent with Fig. 3B, the mixture of two Gaussian components in TW1 and TW2 suggests that most helicase-loading intermediates transitioned from a low-*E*_FRET_ DNA-bent state to a high-*E*_FRET_ DNA-deposited state between TW1 and TW2.

**Figure 4:**
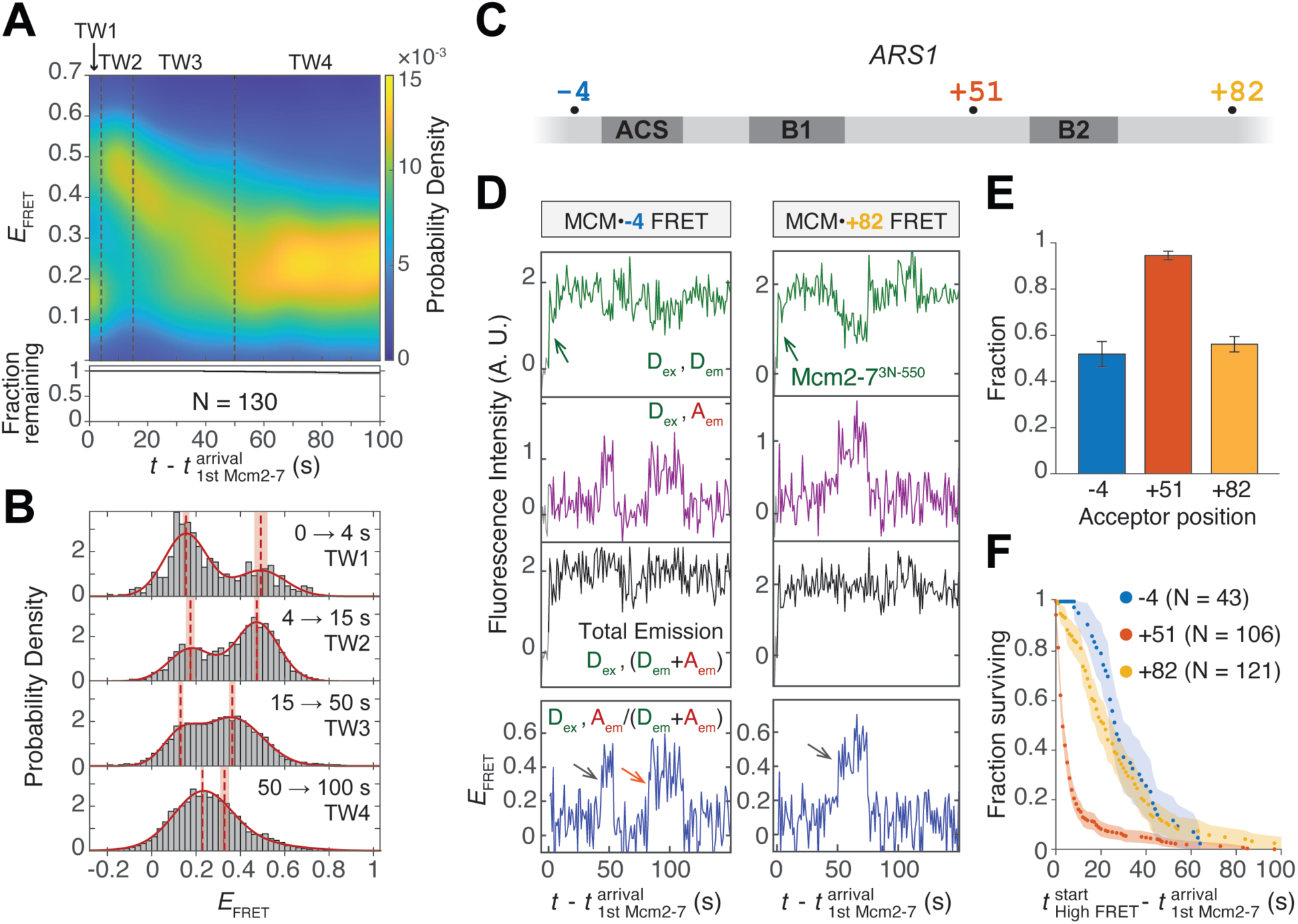
Single Mcm2-7 hexamers slide on DNA after DNA insertion into the central channel. **A.** Heat map showing MCM•+51 *E*_FRET_ distribution changing with time after first Mcm2-7 arrival (see Methods). Data were from 130 salt-stable helicase loading events taken from experiments described in Fig. 3A. The *E*_FRET_ probability density (color scale) was calculated using Mcm2-7 molecules still visible at each time point. The fraction of Mcm2-7 that remain bound (black curve) and the 95% CI (black shading) are shown in the bottom plot. Because all Mcm2-7 molecules represent successfully loaded helicases, the decrease in fraction bound Mcm2-7 was due to termination of experiment rather than dissociation. The heat map is divided into four time windows (TW) as defined by the dashed line boundaries. TW1: 0 - 4s after the first Mcm2-7 arrival; TW2: 4 - 15 s; TW3: 15 - 50 s; and TW4: 50 - 100 s. Time resolution = 1 s. **B.** MCM•+51 *E*_FRET_ in each TW plotted with bin width 0.02. The *E*_FRET_ distributions of each TW were separately fit to two-component Gaussian mixture models (red curves, *SI Appendix*, Table S1). The center values of each Gaussian component (red dashed lines) and the standard errors of these values (red shadings) are shown. **C.** Diagram of *ARS1* origin showing the acceptor modification sites used to detect sliding in the MCM•-4 and MCM•+82 FRET experiments. The +51 site of modification used in the MCM•+51 FRET experiment to detect DNA deposition and sliding is also indicated. **D.** Representative intensity records from FRET experiments with Mcm2-7^3N-550^ and DNA-coupled acceptors at -4 (left) or +82 (right). Green arrow: Mcm2-7^3N-550^ association; grey arrows: the first *E*_FRET_ peak observed; orange arrow: second *E*_FRET_ peak. **E.** Fraction of DNA-bound Mcm2-7 that exhibited at least one high FRET peak in the MCM•-4 FRET, MCM•+51 FRET, and MCM•+82 FRET experiments. Error bars: standard error. **F.** Cumulative distribution of time intervals separating the first Mcm2-7 arrival and the first observation of high *E*_FRET_, including only Mcm2-7 colocalization events that showed high *E*_FRET_. Time resolution of the MCM•+51 experiment (red) is 1 s to allow for direct comparison with the other experiments. Shaded areas: 95% CI; N: the number of Mcm2-7 colocalization events.

After DNA deposition, *E*_FRET_ values decreased in TW3 to a lower level maintained in TW4 (Fig. 4A). Although TW3 shows a gradual decrease in the aggregate heat map (Fig. 4A), single-molecule records show examples of both gradual and sharp *E*_FRET_ decline (*SI Appendix*, Fig. S5). Notably, the peak centers of TW3 and TW4 did not overlap with those of TW1 and TW2 (Fig. 4B), suggesting the presence of new conformational states distinct from the DNA-bent and the DNA-deposited states.

### Mcm2-7 hexamers slide on dsDNA after DNA deposition

Based on previous studies indicating that both Mcm2-7 double hexamers and ORC can slide on DNA (5, 6, 31–33), we hypothesized that the *E*_FRET_ changes after DNA deposition are the result of Mcm2-7 sliding on DNA (*SI Appendix*, Fig. S6A). To confirm Mcm2-7 sliding, we created two additional DNA substrates, each with a single DNA-coupled acceptor at a position located more than 30 bp (> 10 nm) away from the +51 position used to detect DNA deposition (Fig. 4C). The - 4 dye (*ARS1*^-4-Cy5^) is intended to detect leftward Mcm2-7^3N-550^ sliding, and the +82 dye (*ARS1*^+82-Cy5^) rightward sliding (Fig. 4C and *SI Appendix*, Fig. S6B).

In experiments with DNA-coupled acceptors at either the -4 or +82 positions, more than half of the DNA-bound Mcm2-7 molecules exhibited one or more *E*_FRET_ peaks (Fig. 4D-E), implying that the Mcm2-7 molecules can slide leftward or rightward along the DNA. Because some molecules displayed more than one *E*_FRET_ peak (Fig. 4D, orange arrow), we infer that the sliding movement can change direction. For Mcm2-7 colocalization events that displayed high *E*_FRET_, the time to sliding detected by the -4 and +82 dyes is significantly longer than time to deposition after Mcm2-7 arrival (Fig. 4F), which is consistent with sliding occurring only after initial DNA deposition. Together, these findings suggest that after DNA deposition, Mcm2-7 single hexamers can slide back and forth on DNA with no preferential direction.

### Regulation of Mcm2-7 sliding

To determine if Mcm2-7 ATP-hydrolysis impacts sliding, we performed MCM•+51 FRET experiments with six Mcm2-7 ATPase mutants (34), each containing a single R→A mutation in one of the six ATPase arginine-finger motifs present in Mcm2-7. The resulting *E*_FRET_ heat maps show that each of the Mcm2-7 ATPase mutants reached a high *E*_FRET_ state that indicates successful DNA deposition (Fig. 5A). Although we observed a spectrum of different *E*_FRET_ patterns in the mutants (*SI Appendix*, Fig. S7), we consistently observed a prolonged duration of the high *E*_FRET_ state (Fig. 5A) compared to WT Mcm2-7 (Fig. 4A). Thus, efficient Mcm2-7 sliding is initiated by at least one round of ATP hydrolysis at each Mcm2-7 subunit interface.

**Figure 5:**
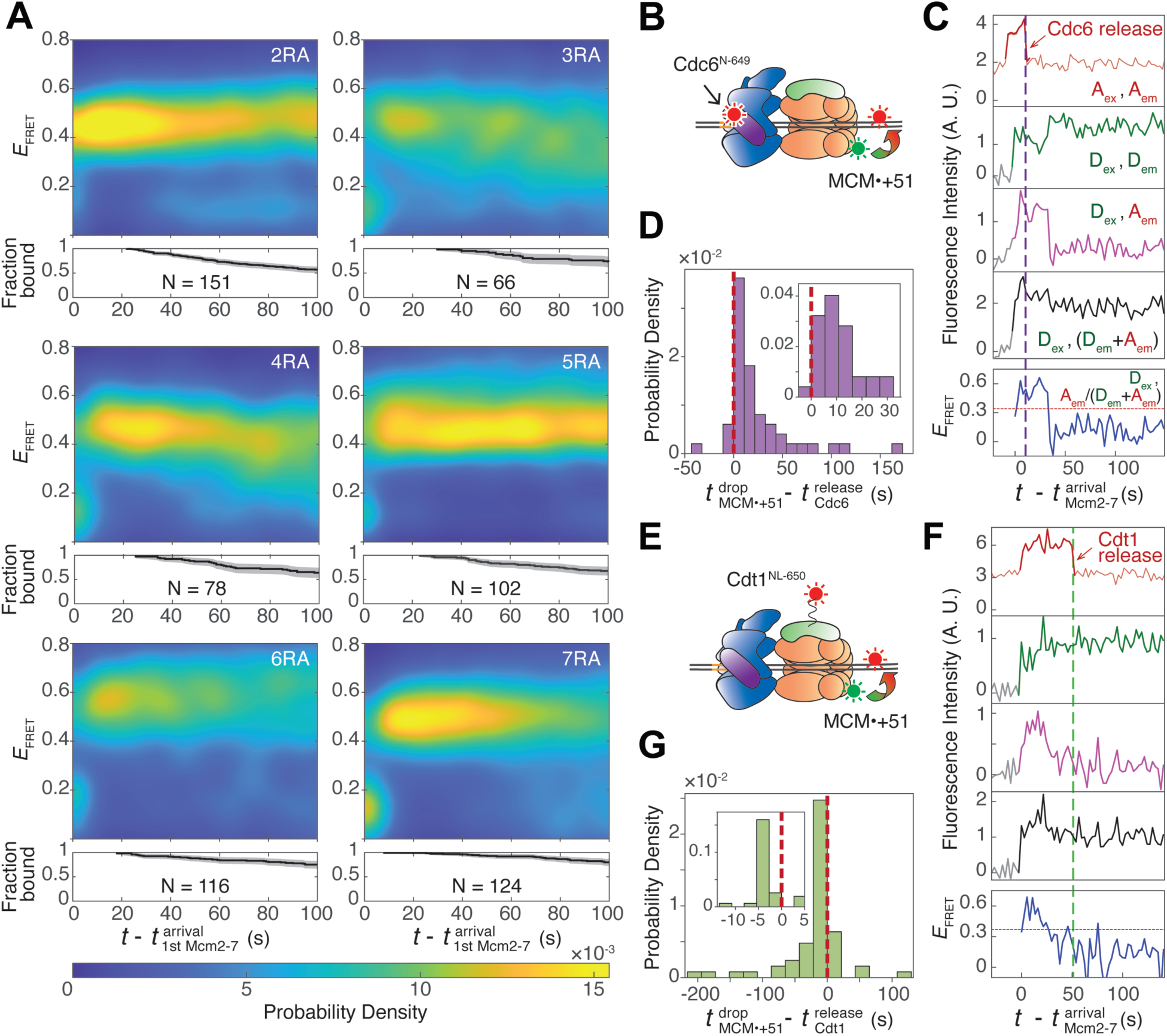
Mcm2-7 sliding requires ATP-hydrolysis and generally begins after Cdc6 release but before Cdt1 release. **A.** MCM•+51 *E*_FRET_ heat maps of each of the six Mcm2-7 arginine-finger ATPase mutants. Plots are as described in Fig. 4A. N: the number of Mcm2-7 colocalization events. Example records are shown in *SI Appendix*, Fig. S7. **B.** Schematic of the MCM•+51 FRET experiment in the presence of Cdc6^N-649^. **C.** Representative record from the experiment in (B). Red arrow and purple dashed line mark Cdc6^N-649^ release. Top plot: acceptor emission during acceptor excitation (A_ex_, A_em_). Fluorescence in light red is contributed by labeled *ARS1*^+51-Cy5^. Darker red interval indicates Cdc6^N-649^ colocalization with DNA. Second plot: donor emission during donor excitation (D_ex_, D_em_). Third plot: acceptor emission during donor excitation (D_ex_, A_em_). Fourth plot: total emission during donor excitation (D_ex_, (D_em_+A_em_)). Bottom plot: effective FRET efficiency (*E*_FRET_) values (D_ex_, A_em_ /(D_em_+A_em_)). Red dotted line marks the threshold (0.34) separating the high and low *E*_FRET_ states. Time resolution = 2.7 s. Additional records are shown in *SI Appendix*, Fig. S8C. **D.** Distribution of time intervals from Cdc6^N-649^ release to the decrease in MCM•+51 *E*_FRET_. A vertical red dashed line marks time interval = 0. Histogram bin width = 10.8 s (4 frames). 42 out of 46 time intervals are positive (Cdc6^N-649^ releases before *E*_FRET_ decrease). Inset: magnified view with bin width = 5.4 (2 frames). **E.** Schematic of the MCM•+51 FRET experiment in the presence of Cdt1^NL-650^. **F.** Representative record from the experiment in (E). Figure descriptions are as (C), except red arrow and green dashed line mark Cdt1^NL-650^ release, and the darker red interval in the top plot indicates Cdt1^NL-650^ colocalization with DNA. Red dotted line marks the threshold (0.37) separating the high and low *E*_FRET_ states. Additional records are shown in *SI Appendix*, Fig. S8F. **G.** Distribution of time intervals from Cdt1^NL-650^ release to the decrease in MCM•+51 *E*_FRET_. A vertical red dashed line marks time interval = 0. Histogram bin width = 21.6 s (8 frames). 46 out of 58 time intervals are negative (Cdt1^NL-650^ releases after *E*_FRET_ decrease). Inset: magnified view with bin width = 2.7 (1 frame).

We next investigated the temporal relationship between the start of Mcm2-7 sliding and the sequential releases of Cdc6 and Cdt1 from the OCCM (Fig. 5B-F). We performed MCM•+51 FRET experiments in the presence of Cdc6 or Cdt1 labeled with a red-excited dye to monitor Cdc6 or Cdt1 release simultaneously with MCM•+51 *E*_FRET_ (Fig. 5B, 5E). We labeled the N-terminus of Cdc6 (Cdc6^N-649^), which is distant from the Mcm3 N-terminal FRET donor based on structural studies (16) and placed the Cdt1-coupled fluorophore on a rigid, N-terminal linker (Cdt1^NL-650^) to separate it from the Mcm3 donor fluorophore (see Methods). Control experiments confirmed that minimal FRET was observed between Mcm2-7^3N-550^ and Cdc6^N-649^ or Cdt1^NL-650^ (*SI Appendix*, Fig. S8). We used alternating red and green laser excitation to monitor protein binding to DNA simultaneously with FRET: green excitation enabled monitoring of FRET and DNA association of green-excited Mcm2-7, while red excitation monitored DNA association of red-excited Cdc6 or Cdt1.

In these MCM•+51 FRET experiments, Cdc6 or Cdt1 release is represented by a decrease in red-excited fluorescence (Fig. 5C and 5F, red arrows). The initiation of Mcm2-7 sliding is indicated by *E*_FRET_ decrease below the defined threshold (Fig. 5C and 5F, red dotted line, *SI Appendix*, Fig. S8B and S8E). More than 90% of DNA-bound Mcm2-7 molecules exhibited *E*_FRET_ decrease only after Cdc6 release (Fig. 5D, positive values, *SI Appendix*, Fig. S8C). In contrast, we found that 80% of Mcm2-7 exhibited the *E*_FRET_ decrease before Cdt1 release (Fig. 5G, negative values, *SI Appendix*, Fig. S8F). Taken together, these findings show that Mcm2-7 sliding primarily occurs after Cdc6 release but before Cdt1 release.

### The initial sliding helicase-loading intermediate includes ORC, Mcm2-7, and Cdt1

Because the initial interactions between ORC and Mcm2-7 remain stable until Cdt1 release (9), our observation that Mcm2-7 sliding typically begins before Cdt1 release strongly suggests that the initial sliding complex also includes Cdt1 and ORC. To confirm this hypothesis and characterize the timing of ORC sliding relative to other steps in helicase loading, we developed a FRET assay that monitors ORC-ACS interaction. We placed a donor fluorophore on the N-terminal face of ORC (ORC^N^, see Methods) and an acceptor fluorophore adjacent to the ACS (*ARS1*^-4-Cy5^) (Fig. 6A). To monitor Mcm2-7 DNA association simultaneously with this ORC^N^•-4 FRET pair, we included Mcm2-7 labeled with a red-excited fluorophore at the N-terminus of Mcm4 (Mcm2-7^4N-650^), which displayed minimal FRET with ORC^N^ due to its long unstructured N-terminal tail (*SI Appendix*, Fig. S9A).

**Figure 6:**
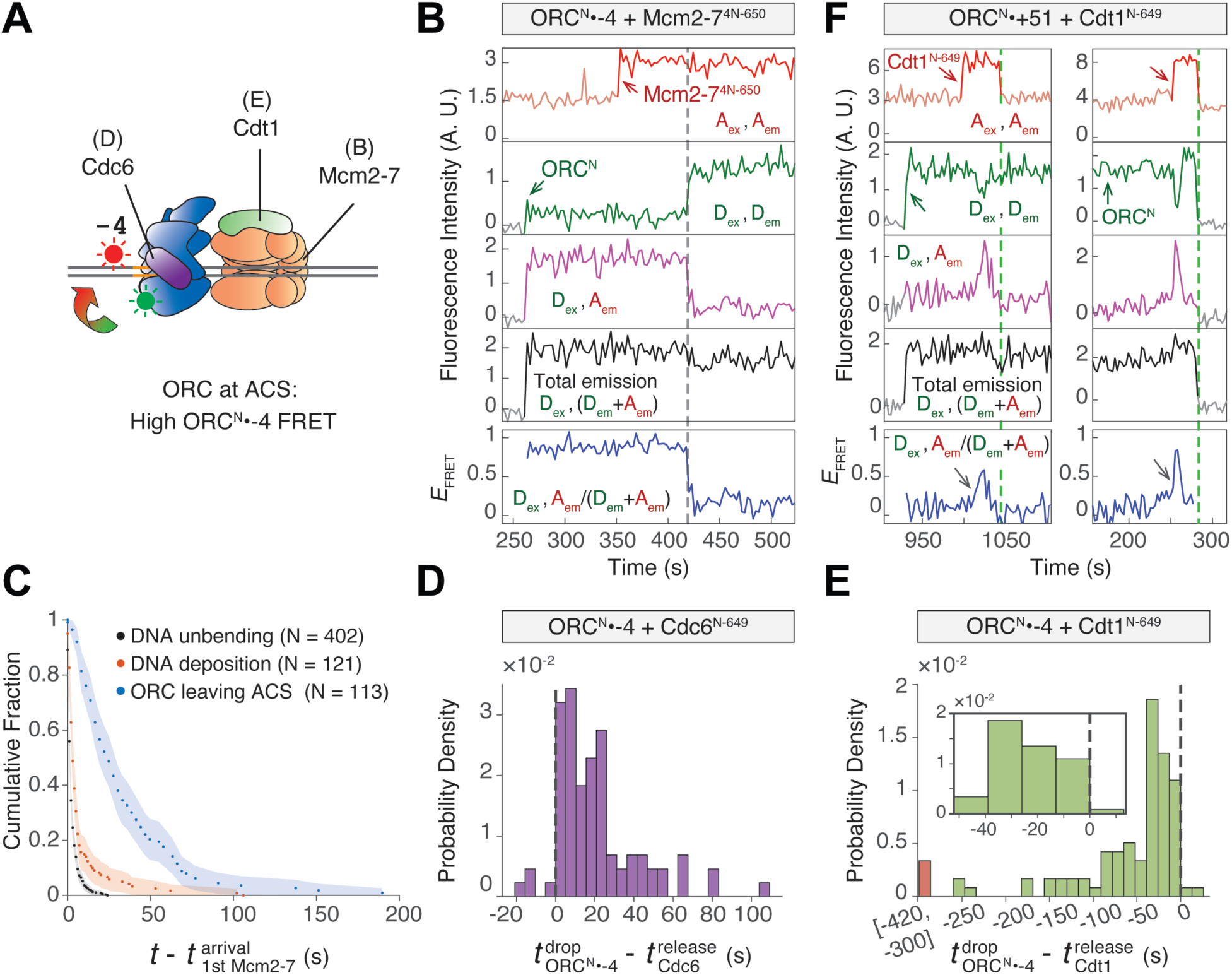
ORC slides away from the ACS as a complex with Mcm2-7 and Cdt1. **A.** Schematic of experiments that monitor ORC-ACS interaction. Black lines indicate the proteins that are labeled with a red-excited fluorophore in the ORC^N^•-4 FRET experiments presented in the corresponding figures. **B.** Example record of the ORC^N^•-4 experiment in the presence of Mcm2-7^4N-550^. Top plot: acceptor emission during acceptor excitation (A_ex_, A_em_). Fluorescence in light red is contributed by DNA. Red arrow and darker red segments of the intensity plot indicate Mcm2-7^4N-650^ colocalization. Second plot: donor emission during donor excitation (D_ex_, D_em_). Green arrow marks ORC^N^ arrival. Third plot: acceptor emission during donor excitation (D_ex_, A_em_). Fourth plot: Total emission during donor excitation (D_ex_, (D_em_+A_em_)). Bottom plot: *E*_FRET_ values (D_ex_, A_em_ /(D_em_+A_em_)). Dashed line marks the decrease of *E*_FRET_, indicating ORC movement away from ACS. More records are shown in *SI Appendix*, Fig. S9B. Time resolution = 2.7 s. **C.** Cumulative distribution for the time interval separating the first Mcm2-7 arrival and one of the following three events: DNA unbending (black), DNA deposition into Mcm2-7 central channel (red), and ORC movement away from ACS (blue). The data from DNA unbending (black) and DNA deposition (red) are from the same experiments as shown in Fig. 3D but with 1 s time resolution to allow direct comparison with the ORC movement data (blue). Shadings: 95% CI; N: number of colocalization events. **D.** Distribution of the time intervals separating ORC^N^•-4 *E*_FRET_ decrease and Cdc6 release, bin width = 5.4 s (2 frames). 80 out of 84 time intervals are positive (*E*_FRET_ decreases after Cdc6 release). Example intensity traces are shown in *SI Appendix*, Fig. S10A. **E.** Distribution of the time intervals separating ORC^N^•-4 *E*_FRET_ decrease and Cdt1 release, bin width = 13.5 s (5 frames). Outliers of values from the range [-420, -300] are grouped in the red bar. Inset: magnified histogram. 89 out of 91 time intervals values are negative (*E*_FRET_ decreases before Cdt1 release). Example records are shown in *SI Appendix*, Fig. S10B. **F.** Two example records of ORC^N^•+51 FRET experiments in the presence of red-labeled Cdt1^N-649^. A schematic of the FRET probes used in this experiment is in *SI Appendix*, Fig. S6C. Grey arrows indicate FRET peaks of ORC sliding to the proximity of the +51 DNA dye. Green dashed line marks Cdt1 release. More records are shown in *SI Appendix*, Fig. S10C.

In this ORC^N^•-4 FRET experiment with labeled Mcm2-7, we observed high *E*_FRET_ before Mcm2-7 arrival, indicating stable ORC binding at the ACS prior to Mcm2-7 recruitment (Fig. 6B, *SI Appendix*, Fig. S9B). We consistently observe the loss of high ORC^N^•-4 *E*_FRET_ after Mcm2-7 arrival (Fig. 6B, dashed line, *SI Appendix*, Fig. S9B), indicating ORC displacement from the ACS. Importantly, we consistently detected a decrease in ORC^N^•-4 FRET for events that led to salt-stable double hexamer formation, indicating that ORC displacement represents an intermediate step on pathway to successful helicase loading (*SI Appendix*, Fig. S9B). By determining the time interval between Mcm2-7 arrival and ORC^N^•-4 *E*_FRET_ decrease, we found that ORC leaving the ACS occurs significantly after DNA unbending or deposition into the Mcm2-7 central channel (Fig. 6C).

The decrease in ORC^N^•-4 *E*_FRET_ could be due to ORC sliding away from ACS or ORC flipping associated with MO formation. If ORC begins sliding simultaneously with Mcm2-7, ORC^N^•-4 *E*_FRET_ decrease would fall between Cdc6 and Cdt1 release (Fig. 5B-G). However, if the *E*_FRET_ decrease corresponds to ORC flipping, which occurs after Cdt1 release (9), we would expect ORC^N^•-4 *E*_FRET_ decrease to follow Cdt1 release. To distinguish between these two possibilities, we determined when ORC leaves the ACS relative to Cdc6 and to Cdt1 release. To this end, we performed ORC^N^•-4 *E*_FRET_ experiments in the presence of red-excited Cdc6^N-649^ or Cdt1^N-649^. Consistent with the predicted distances from structural studies (16), neither Cdc6^N-649^ nor Cdt1^N-649^ exhibits significant *E*_FRET_ to ORC^N^ (*SI Appendix*, Fig. S9A).

ORC^N^•-4 *E*_FRET_ consistently decreased after Cdc6 release (80/84 positive values in Fig. 6D, *SI Appendix*, Fig. S10A). In contrast, ORC^N^•-4 *E*_FRET_ almost always decreased before Cdt1 release (89/91 negative values in Fig. 6E, *SI Appendix*, Fig. S10B). These distributions show that ORC movement away from ACS predominantly begins between Cdc6 release and Cdt1 release, matching the time interval during which Mcm2-7 starts sliding. Importantly, we detected negligible background sliding of ORC in the absence of other helicase-loading factors (*SI Appendix*, Fig. S2), consistent with ORC sliding being temporally controlled during helicase loading.

To detect ORC sliding, we examined *E*_FRET_ between ORC and a DNA probe distant from the ACS. By placing the *ARS1* DNA acceptor at the +51 position, a high ORC^N^•+51 *E*_FRET_ is only expected if ORC moves to the proximity of the +51 dye (*SI Appendix*, Fig. S6C). Importantly, ORC bound at B2 would place the donor fluorophore too far (> 8.2 nm) from the +51 DNA fluorophore to exhibit high *E*_FRET_ (*SI Appendix*, Fig. S6C). In the presence of labeled Cdt1^N-649^, we observed high ORC^N^•+51 *E*_FRET_ peaks that occurred before Cdt1 release (Fig. 6F, *SI Appendix*, Fig. S10C). These *E*_FRET_ peaks indicate that ORC slides away from the ACS while still bound to the C-terminal face of Mcm2-7 and before flipping to form the MO complex.

Given our previous observation that OC_1_M sliding is bidirectional (Fig. 4E), we would expect ORC to revisit the ACS during sliding. If true, then in the ORC^N^•+51 *E*_FRET_ experimental setup, after an initial decrease in *E*_FRET,_ we would sometimes observe increases that represent ORC re-engagement with the ACS. Indeed, 42.9% of ORC molecules exhibited a return to high ORC^N^•+51 *E*_FRET_ signal after the initiation of sliding (*SI Appendix*, Fig. S11A). Consistent with ORC having low affinity for the ACS under these conditions, these rebinding events were short-lived relative to ORC independently binding the ACS (*SI Appendix*, Fig. S11B). Interestingly, only 3.3% of ORC molecules showed a high *E*_FRET_ signal after Cdt1 release, suggesting that Cdt1 release inhibits ORC rebinding to the ACS (*SI Appendix*, Fig. S11A).

Together our studies show that sliding of ORC and Mcm2-7 on the DNA is temporally controlled. This process is triggered by the release of Cdc6 from the OCCM but does not require Cdt1 release. Because the initial ORC-Mcm2-7 interactions remain stable until Cdt1 release (9), we conclude that the OC_1_M complex, consisting of ORC, Cdt1, and Mcm2-7, exhibits bidirectional sliding on dsDNA. ORC sliding away from the ACS represents a third mechanism to reduce ORC affinity for DNA, preparing it to rapidly release from the DNA upon Cdt1 release and disruption of the initial ORC-Mcm2-7 interface. Together, these events drive ORC release from the ACS and facilitate its subsequent flipping, B2 element binding, and MO complex formation.

## Discussion

Our previous studies showed that a single ORC can mediate loading of both helicases found at licensed origins (7, 9). This process requires the same ORC to sequentially bind two distinct and oppositely-oriented sites on the DNA. This model raises the question of how ORC is released from its initial high-affinity binding site ACS.

Using sm-FRET assays to examine ORC and Mcm2-7 interactions with DNA during recruitment of the first Mcm2-7, our studies revealed a coordinated series of changes that promote ORC site-exchange. We propose an updated helicase loading model that incorporates the findings from our study (Fig. 7). DNA-bound ORC consistently induces DNA bending by simultaneously interacting with the ACS and the bend-proximal region (BPR) (Fig. 1D). Shortly after recruitment of the first Mcm2-7 onto ORC-Cdc6 bound to bent DNA (Fig. 7, 1^st^ Mcm2-7 recruitment), DNA unbending is triggered by interactions formed during OCCM assembly, such as between ORC and the WHDs of Mcm4 and Mcm6 (Fig. 2D). This leads to the rapid DNA deposition into the Mcm2-7 central channel (Fig. 3D, Fig. 7, circled 1). Loss of the ORC-BPR interaction destabilizes ORC on DNA (Fig. 1E), and subsequent Cdc6 release (Fig. 7, circled 2) further weakens ORC-DNA interactions (*SI Appendix*, Fig. S1C). Mcm2-7 ATP hydrolysis activity initiates <u>O</u>RC-<u>C</u>dt<u>1</u>-<u>M</u>cm2-7 (OC_1_M) sliding on DNA (Fig. 5A, Fig. 7, circled 3), which both fully releases ORC from the ACS and enables access to B2 elements located at variable distances from the ACS. The sequential events of DNA unbending, Cdc6 release, and OC_1_M sliding progressively reduce ORC stability on DNA (Fig. 7, grey wedge) to promote ORC dissociation from its strong site, allowing for its subsequent rebinding to the inverted B2 element (Fig. 7, ORC Flip). After ORC flipping, B2-bound ORC in the context of the MO complex recruits a second Mcm2-7 in the inverted orientation, completing helicase loading (Fig. 7, 2^nd^ Mcm2-7 recruitment and DH).

**Figure 7:**
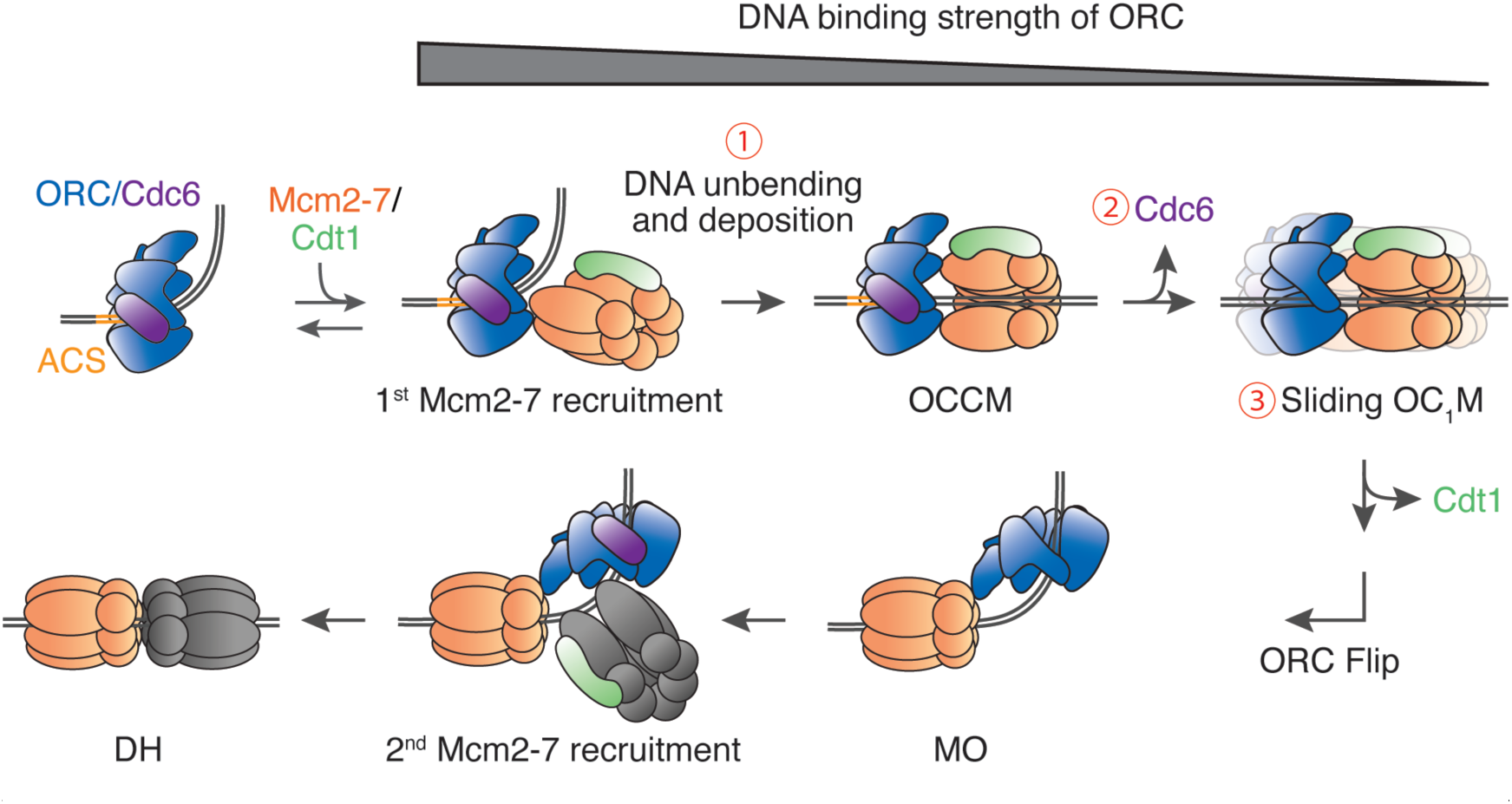
A helicase loading model. We propose that DNA unbending and deposition, Cdc6 release, and OC_1_M sliding (red circled numbers) sequentially lower the DNA binding strength of ORC (grey wedge). This facilitates ORC to release DNA from its central channel and rebind to an inverted binding site (ORC Flip), which enables loading of the second Mcm2-7 in the correct orientation and completes formation of the Mcm2-7 double hexamer (DH).

### Stepwise reduction in ORC stability on DNA

Our finding that ORC is progressively destabilized on DNA is consistent with previous structural studies. ORC makes numerous contacts with the BPR when the DNA is in a bent state (14) and we show that mutation of the amino acids responsible for a subset of these interactions significantly reduces ORC-DNA stability (*SI Appendix*, Fig. S1D). Similar ORC-BPR interactions observed in metazoan ORC suggests that DNA bending is a common mechanism for regulating ORC-DNA complex stability (35). A role for the Mcm4 and Mcm6 WHDs in triggering DNA unbending is consistent with structural studies showing that these domains orient the Mcm2/5 gate next to the BPR for subsequent deposition (36). Notably, the Orc1 basic patch region that interacts with the ACS in the ORC-ACS structure becomes disordered in the OCCM structure, which may further contribute to the lowered ACS affinity of ORC (14, 16).

Following DNA unbending, Cdc6 release serves a dual purpose. Structural studies of the ORC-Cdc6-DNA complex show that Cdc6 binding to ORC completes an ORC-Cdc6 protein ring around the DNA (16, 27). Cdc6 dissociation is thus required to reopen the ORC ring for DNA release from its central channel during ORC flipping. Moreover, the WHD and initiator-specific motif (ISM) of Cdc6 form extensive specific contacts with the ACS (27). Both of these functions are consistent with previous data showing that Cdc6 enhances ORC binding to the ACS (37) and our data that Cdc6 stabilizes DNA-bound ORC (*SI Appendix*, Fig. S1C). The decreased affinity of ORC for the ACS after Cdc6 release would lower the energy barrier for OC_1_M sliding on DNA, which would further reduce specific ORC-DNA interactions. Even upon re-encountering the ACS site during sliding, ORC in this destabilized state fails to maintain stable interactions with the ACS, resulting in its rapid disengagement from the site (*SI Appendix*, Fig. S11). The combined effects of DNA unbending, Cdc6 release, and OC_1_M sliding promote ORC dissociation from DNA necessary for ORC flipping.

Our studies also provide insights into the importance of the ordered release of Cdc6 and Cdt1 from the OCCM. Cdt1 release is associated with the disruption of the interaction between ORC and the C-terminal domain of Mcm2-7 involved in initial Mcm2-7 recruitment (9). Thus, the retention of Cdt1 as sliding commences means that both ORC and Mcm2-7 remain as a complex during initial sliding. Once ORC associates with non-specific DNA, we predict that the stability of ORC on DNA is highly dependent on its interaction with the C-terminal domain of Mcm2-7. When this ORC-Mcm2-7 interaction is disrupted during Cdt1 release, ORC is poised to rapidly dissociate from DNA and seek a new binding site.

Two key questions remain to be addressed by future studies. First, how does ORC remain tethered to the first Mcm2-7 after disengaging from the Mcm2-7 C-terminal domains and DNA? One possibility is that ORC is tethered to the N-terminal domain of Mcm2-7 via Orc6, potentially using the same Orc6-Mcm2 interaction observed in the MO complex (8). Given the long unstructured domain between the Orc6 N-terminal domain and the rest of ORC (26), such an interaction would allow ORC to identify a second binding site without releasing into solution. Second, what causes ORC to preferentially bind to a secondary site in the inverted orientation? Biochemical studies have found that Cdt1 binding to Mcm2-7 relieves the autoinhibitory activity of Mcm6 WHD that blocks Mcm2-7 binding to ORC (38). It is likely that Cdt1 release before MO formation reverts the Mcm6 WHD to its autoinhibitory state. This change would prevent ORC re-association with the Mcm2-7 C-terminus and reduce the likelihood of ORC rebinding to the ACS. Consistent with this hypothesis, we see that release of Cdt1 from the OC_1_M complex significantly reduces ORC rebinding to the ACS (*SI Appendix*, Fig. S11A). Future experiments will be necessary to determine the precise mechanism of ORC binding to the inverted B2 site.

### Energy requirements that lead to ORC flipping

Because the DNA binding strength of ORC is sequentially decreased to facilitate ORC flip and MO formation (Fig. 7, grey wedge), we propose that the resulting increase in the free energy of ORC-DNA interactions is compensated by additional stabilizing interactions or by energy derived from ATP hydrolysis. First, destabilization caused by the loss of ORC-BPR interaction during DNA unbending is offset by stabilizing interactions formed between DNA and the central channel of Mcm2-7 during DNA deposition (MCM-BPR in Fig. 1, Fig. 3), as well as interactions between ORC and Mcm2-7 that involve Mcm4 and Mcm6 WHDs (Fig. 2D). Second, ATP hydrolysis by Cdc6 has been reported to drive Cdc6 release (39). Third, consistent with previous studies showing that Mcm2-7 ATP hydrolysis is required for helicase loading (34, 40), our study demonstrated that Mcm2-7 ATP hydrolysis is required for the onset of OC_1_M sliding (Fig. 5A). A likely explanation for this requirement is the ATP control of a transition of the Mcm2-7 β-hairpin loops from a DNA-engaged (as seen in the OCCM) (3, 16) to a weakly DNA-bound open conformation (as seen in the double hexamer) that favors DNA sliding (2, 41).

Although ATP hydrolysis is important for initiating sliding, our data suggest that OC_1_M movement involves passive diffusion instead of ATP-powered translocation. We observed evidence for movement in both directions from the initial ORC-Mcm2-7 binding sites (Fig. 4D-E), which differs from the biased random walk exhibited by the CMG helicase during active translocation along ssDNA (41). In addition, ORC, single or double Mcm2-7 hexamers, and potential loading intermediates including both ORC and Mcm2-7 have been shown to passively slide on dsDNA (5, 6, 8, 33, 42, 43). Analogous sliding has been observed in many different protein complexes including MutS (44, 45) and Type III restriction enzymes (46).

### Role of sliding during helicase loading

Due to the variable distance between the ACS and B2 elements in different origins, the first Mcm2-7 hexamer must slide a variable distance in either direction for ORC to gain access to the inverted B2 binding site(s). In the case of *ARS1*, in which the OCCM would occupy ACS and B2 simultaneously, OC_1_M sliding would overcome steric obstruction of B2 by the first Mcm2-7. At other origins, such sliding would similarly provide ORC and the first Mcm2-7 access to one or more potential B2 elements. Our data also agrees with previous studies that single roadblocks on either side of the origin do not impact helicase loading (47) because the OC_1_M complex is able to slide away from the roadblocks to access distant B2 elements. The lack of sliding directionality is also consistent with observations that artificial origins with B2 positioned on either side of ACS can support helicase loading (21). A strong requirement for a specific B2 element is observed when flanking roadblocks limit OC_1_M sliding such that only a single B2 element is accessible (47), or in artificial origins where only a single B2-like sequence is present (21). As such, sliding enables ORC to search for another near-ACS sequence in the inverted orientation. Though any ACS-related sequence on either side can serve as a secondary ORC landing site, the distance between the two binding sites is likely confined *in vivo* by origin-flanking nucleosomes that limit sliding (48, 49). As a result, the B2 elements that are found to be important *in vivo* (17, 19) are likely the most accessible secondary sites relative to the ACS.

Our study identifies the OC_1_M as the first sliding intermediate and establishes that sliding is prevented before this intermediate is formed, but it is likely that later helicase-loading intermediates also slide on DNA to search for a stable secondary binding site. Because previous studies proposed that ORC remains tethered to the helicase to prevent its dissociation while exchanging binding sites (8, 9), it is possible that the DNA-bound Mcm2-7 single hexamers travel along DNA while the tethered ORC actively searches for a suitable landing site. Moreover, given that ORC can search DNA via diffusion (32, 33), and the MO complex has been found to occupy DNA sites other than the initial recruitment site (8), the MO complex can likely diffuse on DNA prior to ORC identifying a stable binding site in the inverted orientation. Future experiments will be required to demonstrate and characterize sliding of other helicase-loading intermediates.

## Materials and Methods

### Nomenclature for fluorescently-modified proteins and DNAs

We use a superscript notation to describe proteins and DNA with varying modifications. The superscript notation consists of two parts separated by a dash: the site of modification and the type of modification. If the type of modification is denoted by a number, it refers to the Dylight™ fluorophore conjugated. For example, *ARS1*^+51-BHQ2^ refers to ARS1 origin labeled at the +51 nucleotide position relative to ACS with a Black Hole Quencher-2; Mcm2-7^4N-550^ refers to Mcm2-7 labeled with Dylight 550 at the Mcm4 N-terminus. We used abbreviations for two specific ORC constructs: ORC^C^ for ORC labeled at the C-terminal face with Dylight 550 and ORC^N^ for ORC labeled at the N-terminal face with Dylight 550 (see subsections below).

*SI Appendix*, Table S2 summarizes all the fluorescently-labeled proteins and DNA used this study.

### Preparation of internally-modified DNA (*ARS1*^+51-Cy5^, *ARS1*^+51-BHQ2^, *ARS1*^-4-Cy5^, *ARS1*^+82-Cy5^)

Internally modified *ARS1* DNAs were assembled by ligating two overlapping PCR fragments. A list of oligos (Integrated DNA Technologies) used in this study is provided in *SI Appendix*, Table S3. The internal Cy5 (iCy5) and internal BHQ2 (iBHQ2) fluorophores were directly attached through the DNA phosphate backbone. The biotinylated PCR fragments were generated with 5’ biotin labeled oligos, and the dye-coupled PCR fragments were generated using internally labeled oligos that anneal to the overlap region. The two PCR fragments were ligated as described previously (47) to yield the final 1.4 kb DNA construct containing a biotin end modification and an internal dye modification around the point of ligation.

### Protein purification

Wild-type (WT) Mcm2-7 and WT ORC were purified as described previously (34). WT Cdc6 and Cdc6^N-649^ were purified as described (7, 29).

### Preparation of ORC^N^

An *S. cerevisiae* strain co-expressing all codon-optimized ORC subunits (50) was used to prepare internally labeled ORC^N^. A 3xFLAG tag was inserted at the ORC1 N-terminus, and an s6 tag (GDSLSWLLRLLN) replaced Orc1 residues 314-325, an unstructured region of Orc1 (2, 14–16). ORC was purified using FLAG affinity resin (Sigma-Aldrich) as previously described (9). ORC, SFP synthase (NEB), and Dy550-CoA (Thermo Scientific) were incubated at a 1:3:30 ratio for 30 min at room temperature. Labeled ORC was then purified on a Superdex 200 Increase 10/300 gel filtration column.

### Sortase-mediated protein labeling

Fluorescence labeling at the N or C terminus of proteins were performed via sortase-mediated coupling as described (9).

### Preparation of ORC^C^ and ORC^C-BPR^

*S. cerevisiae* strains overexpressing the codon-optimized ORC subunits with the indicated mutations were grown, arrested, induced, harvested, and lysed as described above. Following elution off the FLAG resin, peak fractions were coupled to Dylight 550 via sortase as above.

### Preparation of Mcm2-7^3N-550^ (WT and RA mutants) and Mcm2-7^4N-650^ (WT and WHD mutants)

*S. cerevisiae* strains co-expressing all Mcm2-7 subunits with the appropriate modifications were grown, arrested, induced, and harvested. Mcm2-7 protein was purified using FLAG resin and labeled via sortase-mediated conjugation as above.

For the Mcm2-7 WHD mutants, 4ΔWHD is a C-terminal truncation mutant lacking residues 854-933; 6ΔWHD lacks residues 839-1017; 4Δ6ΔWHD combines both truncations.

### Preparation of Cdc6^N-649^, Cdt1^N-649^, and Cdt1^NL-650^

Cdc6^N-649^ and Cdt1^N-649^ was purified as described (7). To separate the Cdt1-conjugated fluorophore from the donor fluorophore on Mcm2-7^3N-550^, a 3x FLAG tag and a rigid alpha-helical (EAAAK)_10_ linker (51) was placed at the N-terminus of Cdt1 after the sortase-recognition motif GGG. Cdt1 was purified via FLAG resin and labeled at the N-terminus via sortase-mediated conjugation as described above.

### Single-molecule assays

A micro-mirror total internal reflection microscope was used to perform multiwavelength single-molecule imaging (24). Glass slides were functionalized with PEG and Biotin-PEG; fiducial markers and biotinylated DNA were coupled to the slide as described (9). All reactions were performed in buffer containing 25 mM HEPES-KOH pH 7.6, 0.3 M potassium glutamate, 5 mM Mg(OAc)_2_, 3 mM ATP, 1 mM dithiothreitol, 1 mg/ml bovine serum albumin, with an oxygen scavenging system (glucose oxidase/catalase) and 2mM Trolox (52). ORC^C^•+51 and ORC^N^•-4 experiments contained 1 nM ORC, 1 µM 60 bp nonspecific DNA generated by annealing two oligonucleotides (*SI Appendix*, Table S3), and 3 nM Cdc6 if specified. 5-10 nM Mcm2-7/Cdt1 was added for helicase loading assays. DNA molecules labeled with Alexa488 or Cy5 were identified before the experiments using 488 nm or 633 nm excitation respectively. Three different protocols were used for experimental acquisition: 1) Reactions containing only green-excited proteins: continuous 532 nm excitation of specified exposure time; 2) Reactions containing green-excited ORC, red-excited Mcm2-7, and quencher-labeled DNA: simultaneous 532 nm and 633 nm excitation; 3) Reactions containing green-excited ORC, red-excited Mcm2-7, and red-excited DNA: alternating 532 nm and 633 nm excitation. Cy5 internally-labeled DNA that did not photobleach were identified by 633 nm excitation after the experiments. To identify salt-resistant helicases, a high-salt buffer containing 25 mM HEPES-KOH pH 7.6, 0.5 M KCl, 5 mM Mg(OAc)_2_ was applied to wash away loading intermediates at the end of the experiments.

### FRET data analysis

Data was analyzed as described previously (7) and fluorescence intensity values were corrected for background fluorescence as described (53). Apparent FRET efficiency (*E*_FRET_) calculations were performed as described (9).

*E*_FRET_ values were maximum likelihood fit using the models and yielding the fit parameters given in *SI Appendix*, Table S1. The *E*_FRET_ threshold used to differentiate the two *E*_FRET_ states was defined as the position of the trough in the fit. An *E*_FRET_ transition was considered to have taken place once the *E*_FRET_ value crossed the threshold for at least two consecutive frames.

### ORC apparent dissociation rate in high and low *E*_FRET_ states

In the ORC^C^•+51 FRET experiment, we used the threshold 0.28 to distinguish between the high and low *E*_FRET_ states. The frequency of ORC release (Fig. 1E) in its high and low *E*_FRET_ states was determined by first identifying the *E*_FRET_ state of ORC in the current frame n. The next frame (frame n+1) was then assessed to determine whether ORC was released into solution, which is indicated by the decrease of green-excited fluorescence back to background levels. The frequency of release was calculated by dividing the number of ORC dissociation events by the total number of frames in the corresponding *E*_FRET_ state.

### *E*_FRET_ heat maps

The *E*_FRET_ heat maps in Fig. 4A and 5A were generated using MATLAB code adapted from https://github.com/gelles-brandeis/jganalyze. The code used two-dimensional kernel density estimation, with a normal kernel function (standard deviation of time axis = 5 s and *E*_FRET_ axis 0.05) to generate a heat map of *E*_FRET_ values against time (0-100 seconds after first Mcm2-7 arrival). Each vertical slice represents the probability density function of *E*_FRET_ values at the particular time interval. The time axis has a resolution of 1 second to match the time resolution of the datasets, and the *E*_FRET_ axis has a resolution of 0.005. Normalization was performed such that the density estimate at each time slice integrates to one.

### Ensemble OCCM formation assays

Ensemble OCCM formation assays (*SI Appendix*, Fig. S1A) were done as described previously (34).

### Data availability

Source data for the single-molecule experiments are provided as Matlab “intervals” files that can be read and manipulated by the program imscroll (https://github.com/gellesbrandeis/CoSMoS_Analysis). The source data are archived at doi: 10.5281/zenodo.7814499.

## Acknowledgements

This work was supported by NIH grants R01 GM147960 (SPB and JG) and R01 GM81648 (JG). AZ was supported by an Angela Leong Fellowship. SPB is an investigator with the Howard Hughes Medical Institute. This work was supported in part by the Koch Institute Support Grant P30-CA14051 from the NCI. We thank the Koch Institute Swanson Biotechnology Center for technical support.

