## Supplementary_File_1 for "Changing protein-DNA interactions promote ORC binding site exchange during replication origin licensing"

**This PDF file includes:**

Figures S1 to S11
Tables S1 to S3

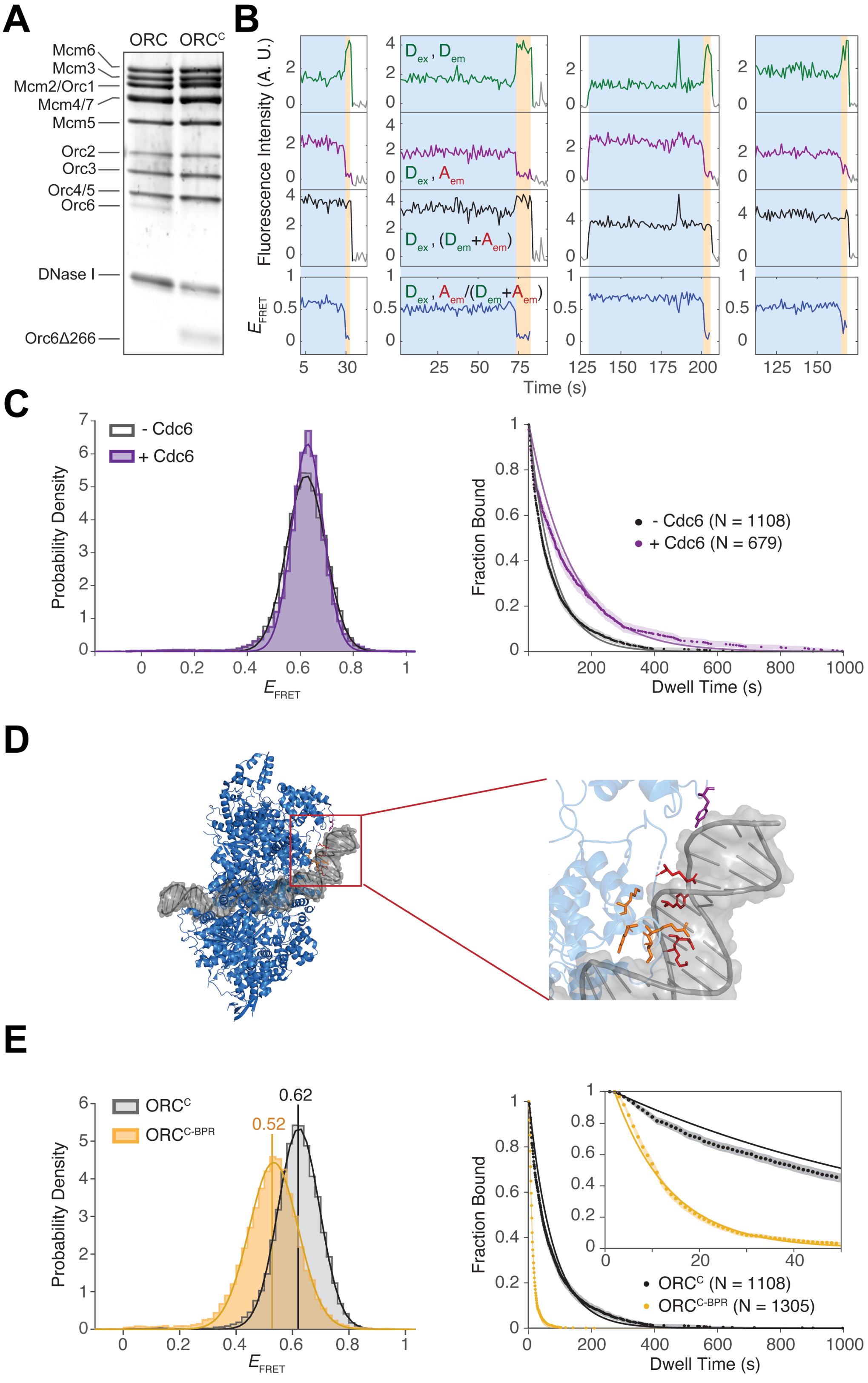


**Figure S1: Additional data on the ORC$^{C}$•+51 experiment in Fig. 1.**

(A) *In vitro* OCCM formation assay showing that ORC$^{C}$ is not defective in OCCM formation compared to WT ORC. The indicated ORC constructs with WT Mcm2-7, Cdc6, and Cdt1 were incubated with *ARS1* DNA in buffer containing ATPγS to capture OCCM complexes. The DNA-bound proteins were released by DNase I treatment and analyzed using SDS-PAGE. The Orc6Δ266 band indicates the truncated Orc6 subunit in the ORC$^{C}$ construct.

(B) Additional records at four DNA molecules from the same ORC$^{C}$•+51 experiment as Fig. 1C. Although ORC-DNA is primarily in the high $E_{\text{FRET}}$ state (blue background), periods of low $E_{\text{FRET}}$ state (yellow background) were occasionally detected before dissociation. Time resolution = 1 s.

(C) Cdc6 does not alter the extent of origin DNA bending but increases ORC dwell time on DNA. Left: distribution of ORC$^{C}$•+51 $E_{\text{FRET}}$ values in the absence (grey) or presence of Cdc6 (purple). Data are fit to one-component Gaussian models (solid curves, *SI Appendix*, Table S1). Right: survival plot of ORC$^{C}$ dwell intervals on DNA in the absence (grey) or presence of Cdc6 (purple). Data are fit to single exponential functions (solid curves, *SI Appendix*, Table S1). Shading: 95% CI; N: the number of ORC molecules colocalized on DNA.

(D) Cryo-EM structure of ORC bound to bent *ARS1* (PDB ID: 5ZR1). The zoomed-in view highlights the 9 residues within the ORC basic patch region directly contacting the phosphate backbone of BPR. These residues are mutated to alanine in the ORC$^{C\text{-}BPR}$ mutant. Orange: Orc2 K251, R254, T255, K258; Red: Orc5 R360, Y363, R366, K367; Purple: Orc6 Y277.

(E) Interaction between ORC and the BPR DNA partially contributes to DNA bending and stabilizes ORC on DNA. Left: distribution of ORC$^{C}$•+51 $E_{\text{FRET}}$ values when ORC$^{C}$ (grey) or ORC$^{C\text{-}BPR}$ (yellow) is colocalized on DNA. Data are fit to one-component Gaussian models (solid curves, *SI Appendix*, Table S1) with centers indicated by vertical lines. Right: survival plot of dwell intervals of ORC$^{C}$ (grey, same as (C)) or ORC$^{C\text{-}BPR}$ (yellow) on DNA. Data are fit to single exponential functions (solid curves, *SI Appendix*, Table S1). Inset: magnified view. Shading: 95% CI; N: the number of ORC molecules colocalized on DNA.

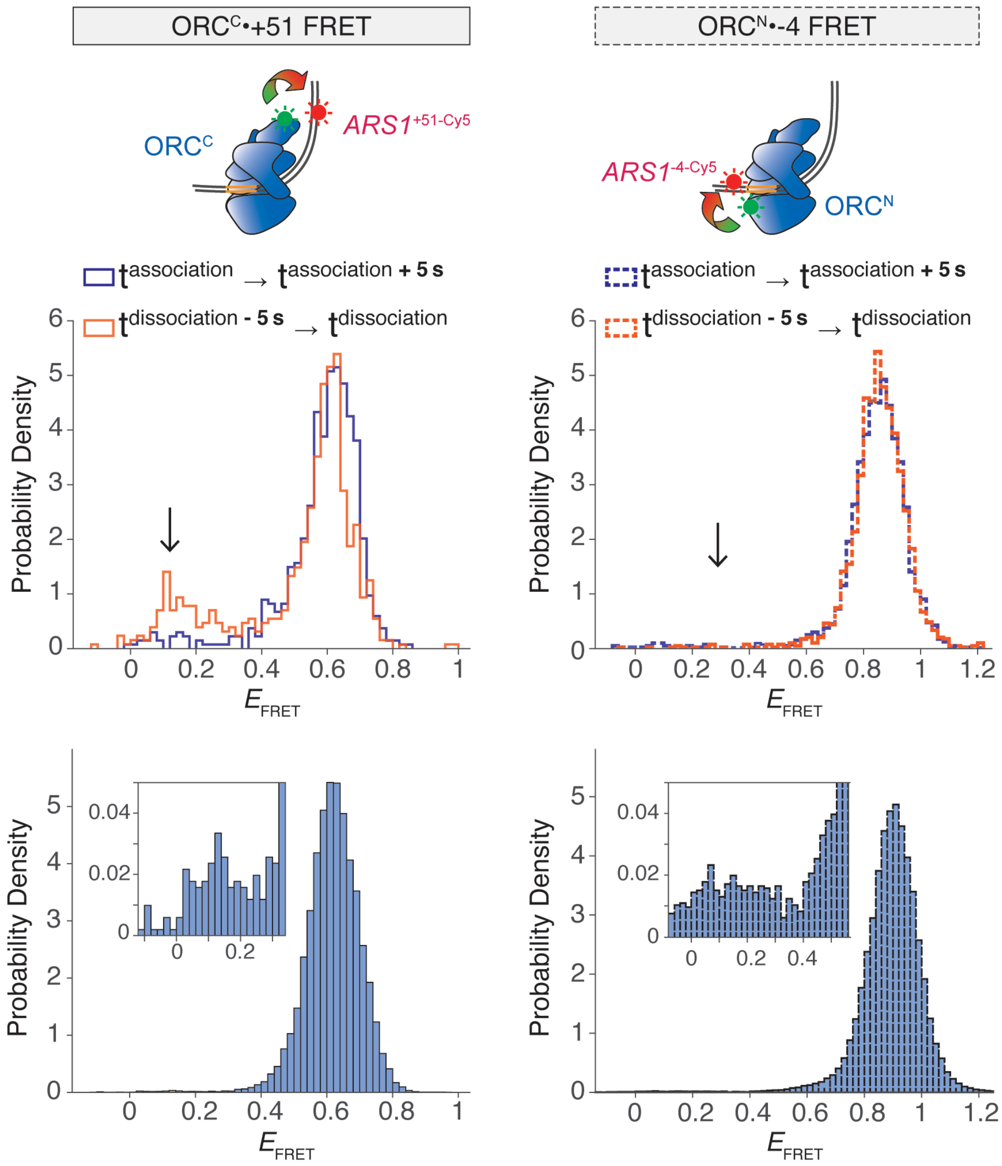


**Figure S2: ORC-ACS interaction is stable while ORC-BPR interaction is occasionally disrupted before ORC dissociation.**

Experiment monitoring DNA bending via interaction between ORC$^{C}$ and *ARS1*$^{+51\text{-}Cy5}$ (left), is compared to experiment monitoring ACS binding via interaction between ORC$^{N}$ and *ARS1*$^{+4\text{-}Cy5}$ (right, more in Fig. 6). For each ORC colocalization event, $E_{FRET}$ values during the first five seconds of colocalization (purple, top) and the last five seconds of colocalization (orange, top) are plotted as histograms. The grey arrows point toward $E_{FRET}$ values in the low state. Compared to ORC$^{C}$•+51 (left), the low ORC$^{N}$•-4 $E_{FRET}$ values (right) were not enriched immediately before ORC dissociation into solution, indicating that ORC remains bound at the ACS but DNA occasionally unbends before dissociation. The bottom panels show the distribution of $E_{FRET}$ values at each frame of ORC colocalization on DNA, and the zoomed-in views in the insets show the distributions of low $E_{FRET}$ values. Number of ORC$^{C}$ colocalization events (left) = 186; number of ORC$^{N}$ colocalization events (right) = 424.

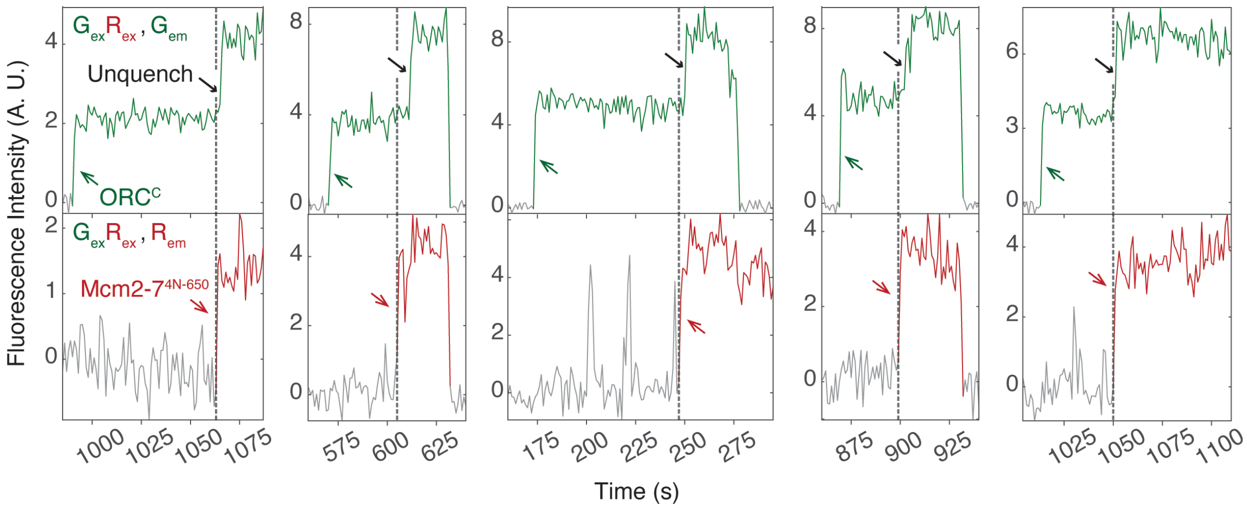


**Figure S3: Additional examples of the ORC•+51 quencher experiment.**

Five additional records from the same experiment as Fig. 2A-B. The vertical dashed line indicates arrival times of the first Mcm2-7. Green, red, and black arrows mark $ORC^{C}$ arrival, Mcm2-$7^{4N\text{-}650}$ arrival, and time of unquenching (DNA unbending) respectively.

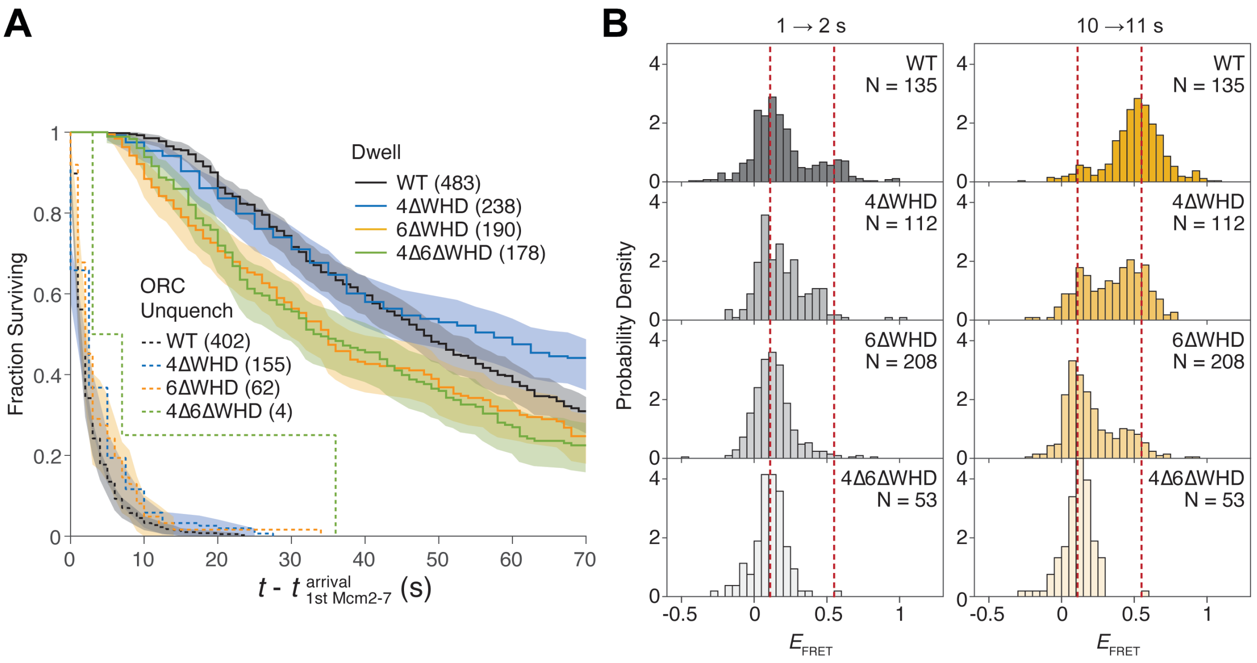


**Figure S4: Mcm4 and Mcm6 WHD mutants exhibit defects in DNA unbending and DNA deposition that are not due to early Mcm2-7 departure.**

(A) DNA unbending defects are not due to early departure of the Mcm2-7 WHD mutants. Cumulative distributions of ORC unquenching (dashed) and Mcm2-7 dwell intervals on DNA (solid) after the first Mcm2-7 arrival. Data are from the $ORC^{C}$•+51 unquenching experiments as in Fig. 2D, which included different labeled Mcm2-7 constructs: WT (black), 4ΔWHD (blue), 6ΔWHD (orange), and 4Δ6ΔWHD (green). Numbers in parentheses represent the number of unquenching or DNA-colocalization events, and shaded areas represent 95% CIs. Not enough ORC unquenching events were observed in the 4Δ6ΔWHD mutant experiment to calculate the CI. Unquenching data from WT Mcm2-7 (black dashed line) are as described in Fig. 2C but collected at 1 s time resolution to allow for direct comparison with the mutant data.

(B) Mcm2-7 WHD mutants that are defective in DNA unbending are also defective in DNA deposition. Comparison of WT and Mcm2-7 WHD mutant MCM•+51 $E_{FRET}$ data using the same experimental scheme as Figure 3A. The plots illustrate MCM•+51 $E_{FRET}$ distributions from 1 - 2 s (left) and 10 - 11 s (right) after first Mcm2-7 arrival. Data shown in the top row is identical to that in Fig. 3C. N indicates the number of Mcm2-7 colocalization events. Red dashed lines correspond to those in Fig. 3C. Transition into the high $E_{FRET}$ state in the 10 - 11 s time window reflects DNA deposition. The 4ΔWHD, 6ΔWHD, and 4Δ6ΔWHD mutants show progressive reduction in DNA deposition, which matches the decreasing fraction of DNA unbending illustrated in Fig. 2D.

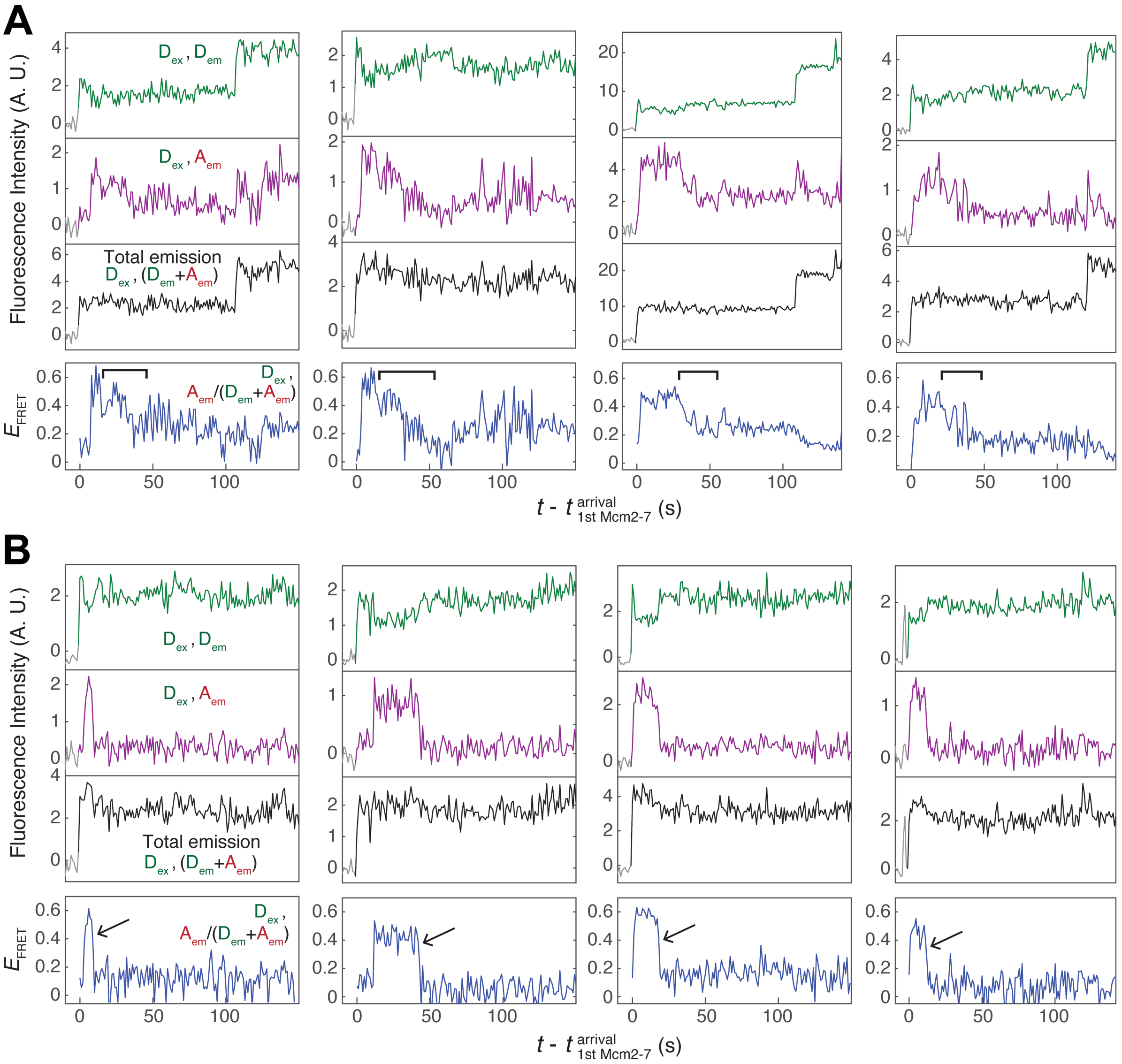


**Figure S5: Decrease in MCM•+51 $E_{FRET}$ can be gradual or abrupt.**

Example records of salt-resistant Mcm2-7$^{3N-550}$ exhibiting MCM•+51 FRET decrease that are (A) gradual, marked by square brackets, or (B) abrupt, marked by arrows. Figure descriptions are as Fig. 3B.

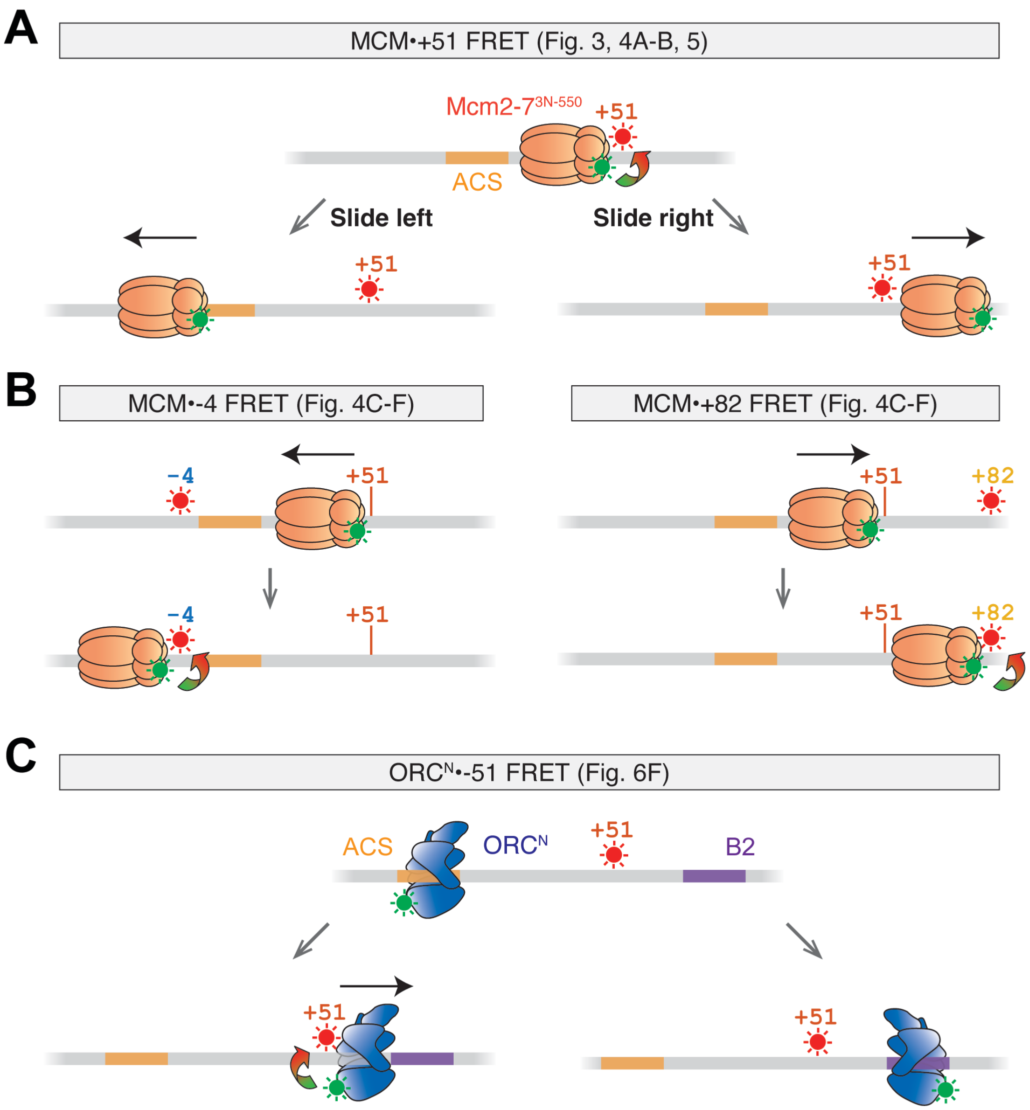


**Figure S6: FRET assays that detect Mcm2-7 or ORC sliding on *ARS1* DNA.**

(A) A red-excited DNA acceptor placed at the DNA +51 position relative to ACS detects DNA deposition. Sliding of Mcm2-7$^{3N-550}$ to either the left or right would separate the Mcm2-7 donor and the DNA acceptor. Note that Mcm2-7 movement to the right involves sliding over the DNA dye.

(B) Mcm2-7$^{3N-550}$ sliding to the left results in MCM•-4 FRET, whereas sliding to the right results in MCM•+82 FRET. Note that Mcm2-7$^{3N-550}$ must slide over the DNA dye to interact with the -4 dye. Vertical orange lines mark the +51 position that is in proximity with the Mcm2-7$^{3N-550}$ donor during initial recruitment.

(C) Left: ORC$^{N}$ sliding from the ACS to the right results in FRET with the +51 DNA dye. Right: A strong FRET is not expected when ORC is bound to B2, as the acceptor dye placed at +51 is estimated to be at least 8.2 nm away from the donor on ORC$^{N}$.

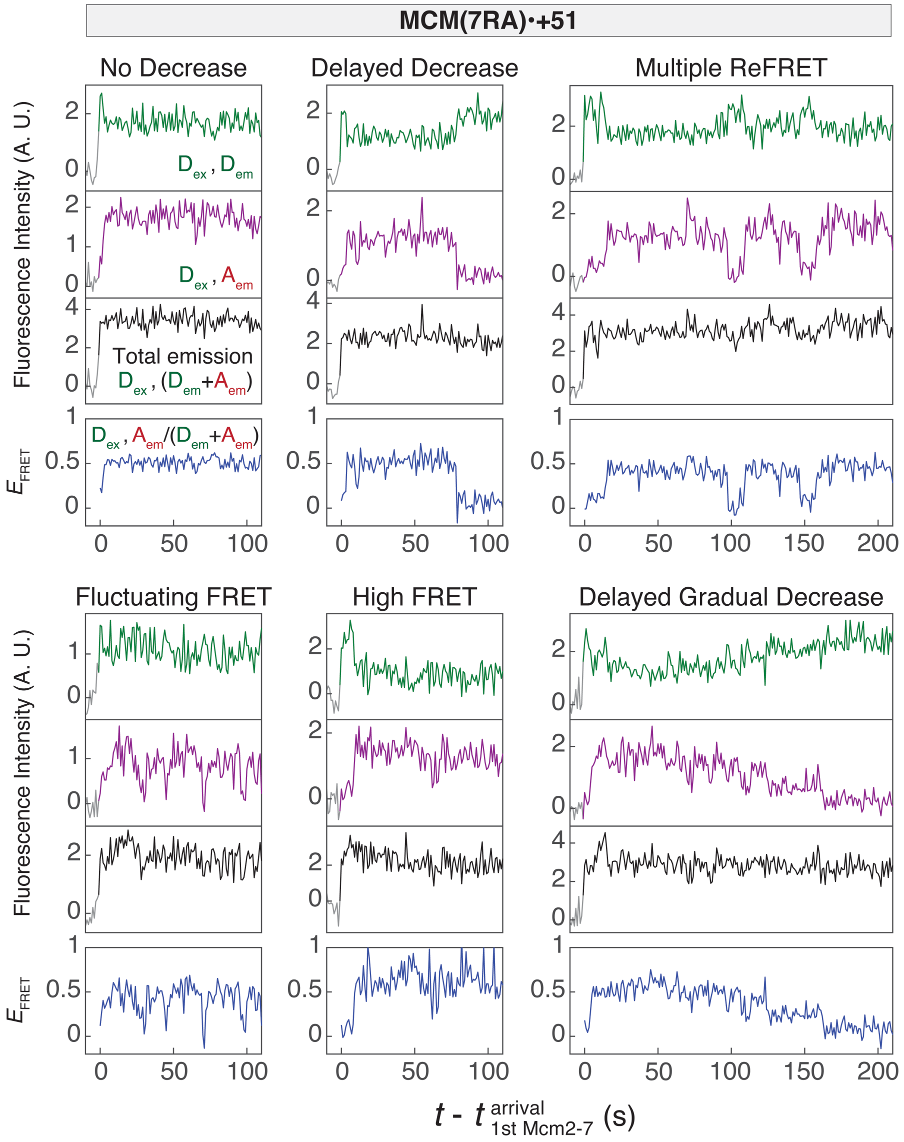


**Figure S7: Mcm2-7 ATPase mutants exhibit a spectrum of MCM•+51 $E_{\mathrm{FRET}}$ patterns but consistently show delayed high-to-low $E_{\mathrm{FRET}}$ transitions.**

Representative single-molecule records from the MCM•+51 $E_{\mathrm{FRET}}$ experiment using the Mcm7RA mutant. The six distinct categories highlight the diverse $E_{\mathrm{FRET}}$ patterns observed. A consistent feature across all categories is the prolonged high $E_{\mathrm{FRET}}$ state.

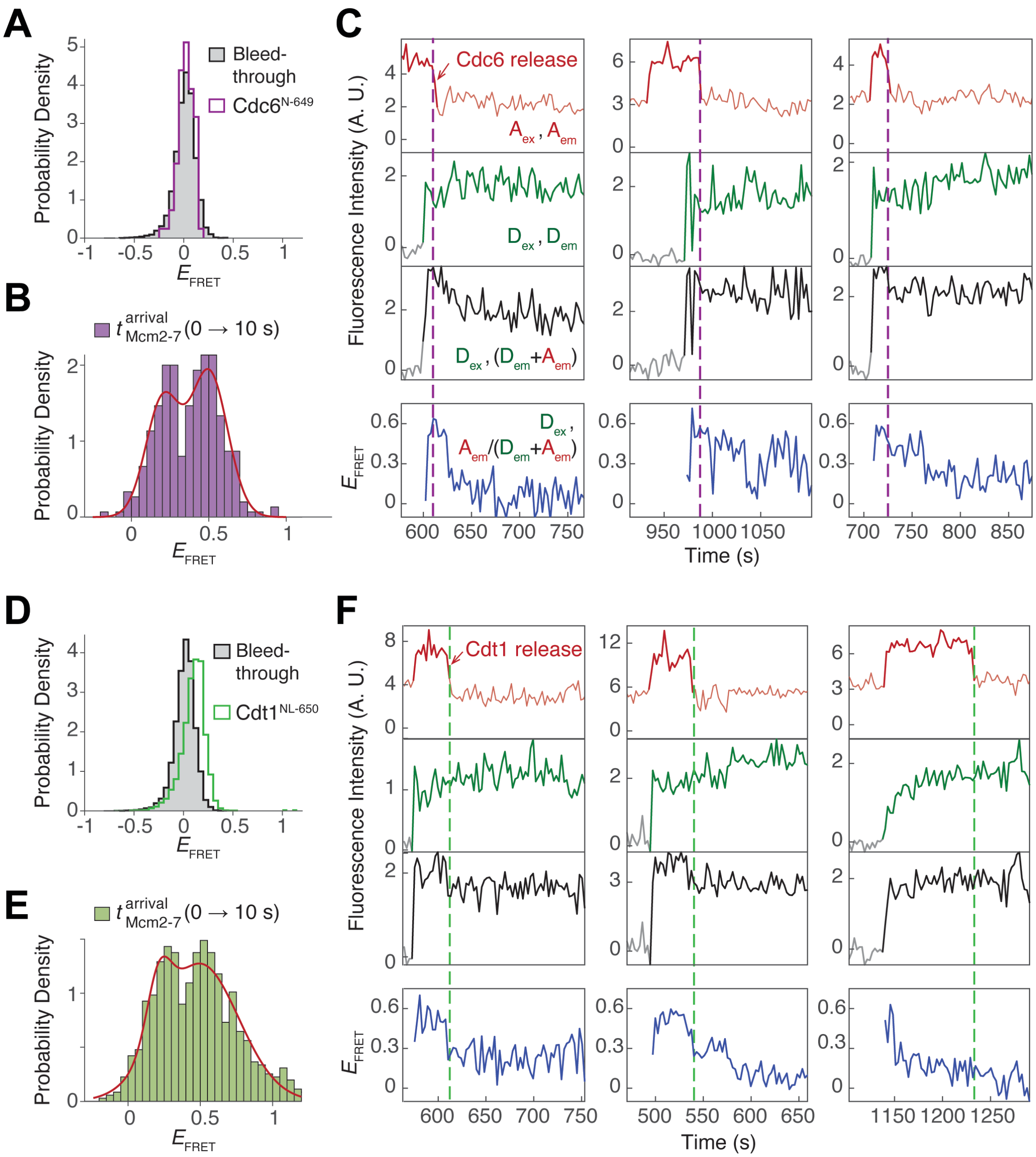


**Figure S8: Labeled Cdc6 and Cdt1 do not significantly interfere with MCM•+51 $E_{FRET}$ measurements**.

(A) Control experiment to assess $E_{FRET}$ between Mcm2-7$^{3N-550}$ and Cdc6$^{N-649}$ in *ARS1* DNA that is not labeled with a FRET acceptor. The grey histogram represents the donor-excited acceptor emission signal from Mcm2-7$^{3N-550}$ colocalization alone. Due to the absence of an acceptor fluorophore, the signal calculated for Mcm2-7$^{3N-550}$ alone represents bleed-through signal. The purple histogram represents the distribution of $E_{FRET}$ during colocalization of both Mcm2-7$^{3N-550}$ and Cdc6$^{N-649}$ on DNA. Similarity of the grey and purple histograms indicates that there is no appreciable FRET signal between Mcm2-7$^{3N-550}$ and Cdc6$^{N-649}$.

(B) Distribution of $E_{FRET}$ values in the first 10 seconds of Mcm2-7 arrival from the MCM•+51 FRET experiment in the presence of Cdc6$^{N-649}$ (as shown in Fig. 5B). Data from 46 Mcm2-7 colocalization events were plotted and fit to a two-component Gaussian model (red curve). Consistent with Fig. 3C and 4B, the low and high Gaussian components represent DNA-bent and DNA-deposited states. The trough value (0.34) from the fit was used as the threshold value to distinguish the low and high $E_{FRET}$ states.

(C) Additional records from the experiment shown in Fig. 5B. Descriptions are as Fig. 5C.

(D) Same as (A) except in an experiment substituting Cdc6$^{N-649}$ with Cdt1$^{NL-650}$ (green). Similarity of the grey and green histograms indicates that there is no appreciable FRET between Mcm2-7$^{3N-550}$ and Cdt1$^{NL-650}$.

(E) Distribution of $E_{FRET}$ values in the first 10 seconds of Mcm2-7 arrival from the MCM•+51 FRET experiment in the presence of Cdt1$^{NL-650}$ (as shown in Fig. 5E). Data from 58 Mcm2-7 colocalization events were plotted and fit to a two-component Gaussian model (red curve). Consistent with Fig. 3C and 4B, the low and high Gaussian components represent DNA-bent and DNA-deposited states. The trough value (0.37) from the fit was used as the threshold value to distinguish the low and high $E_{FRET}$ states.

(F) Additional records from the experiment shown in Fig. 5E. Descriptions are as Fig. 5F.

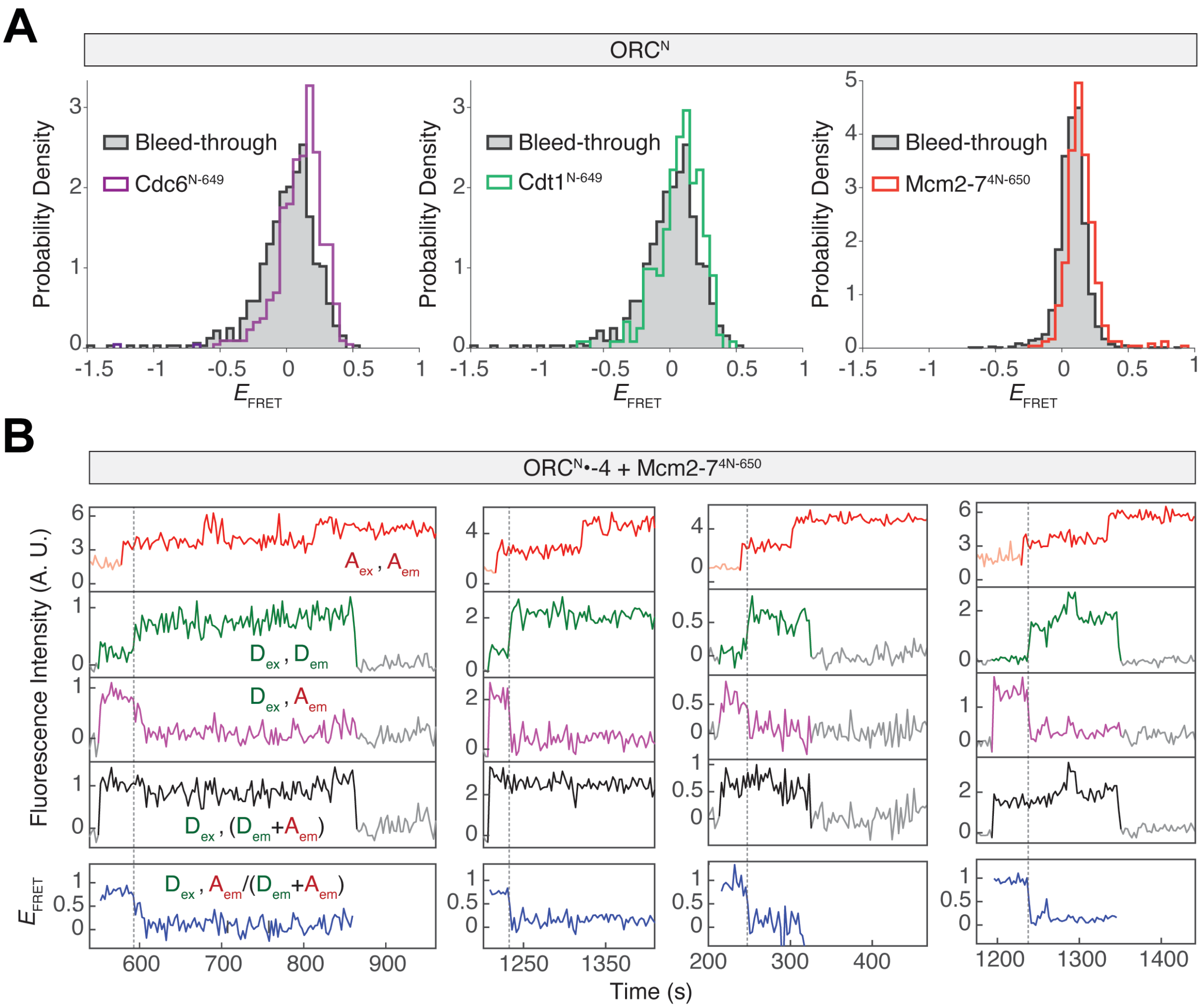


**Figure S9: ORC$^{N}$•-4 $E_{FRET}$ decreases after the first Mcm2-7 arrival and its measurement is not significantly impacted by the presence of other red-excited proteins.**

(A) ORC$^{N}$ show minimal $E_{FRET}$ with three red-excited fluorescent proteins. $E_{FRET}$ distributions of ORC$^{N}$ with Cdc6$^{N-649}$ (purple, left), Cdt1$^{N-649}$ (green, middle), and Mcm2-7$^{4N-650}$ (orange, right) are compared to bleed-through signal of ORC$^{N}$ alone in the donor-excited acceptor emission channel (grey). DNA molecules in these experiments are not labeled with red-excited fluorophores.

(B) Additional records from the experiment in Fig. 6B. Dashed lines: $E_{FRET}$ decrease. All records represent productive helicase-loading events that resulted in salt-stable double hexamers.

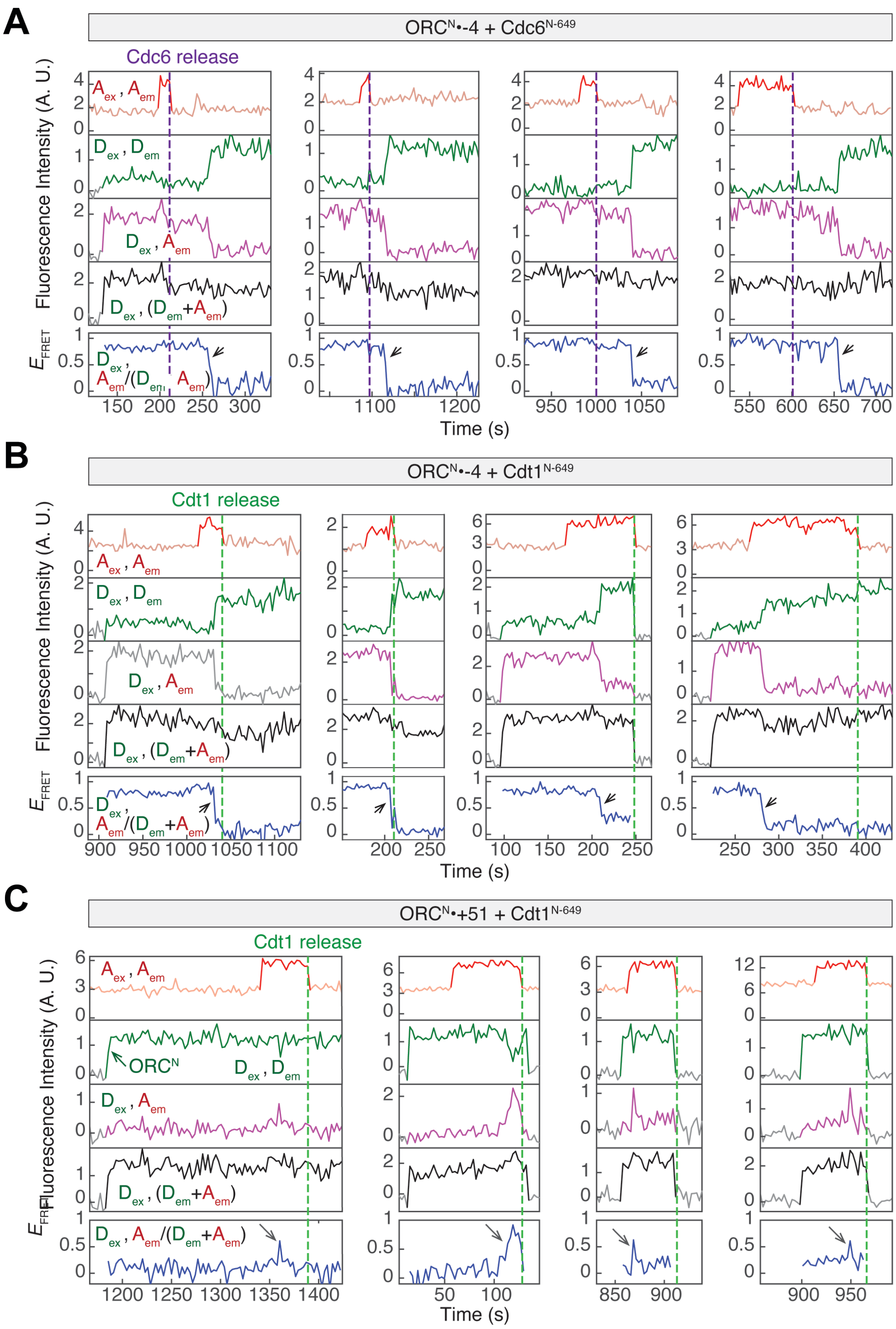


**Figure S10: ORC sliding begins after Cdc6 release but before Cdt1 release.**

Example records of the $ORC^N$•-4 experiment with fluorescently labeled (A) $Cdc6^{N\text{-}649}$ or (B) $Cdt1^{N\text{-}649}$, as described in Fig. 6D-E. Purple dashed line: Cdc6 release; Green dashed line: Cdt1 release; Grey arrows: decrease in $ORC^N$•-4 $E_{FRET}$.

(C) Additional records of the $ORC^N$•+51 FRET experiment as described in Fig. 6F. Schematic of this experiment is shown in *SI Appendix,* Fig. S6C. Green dashed line: Cdt1 release.

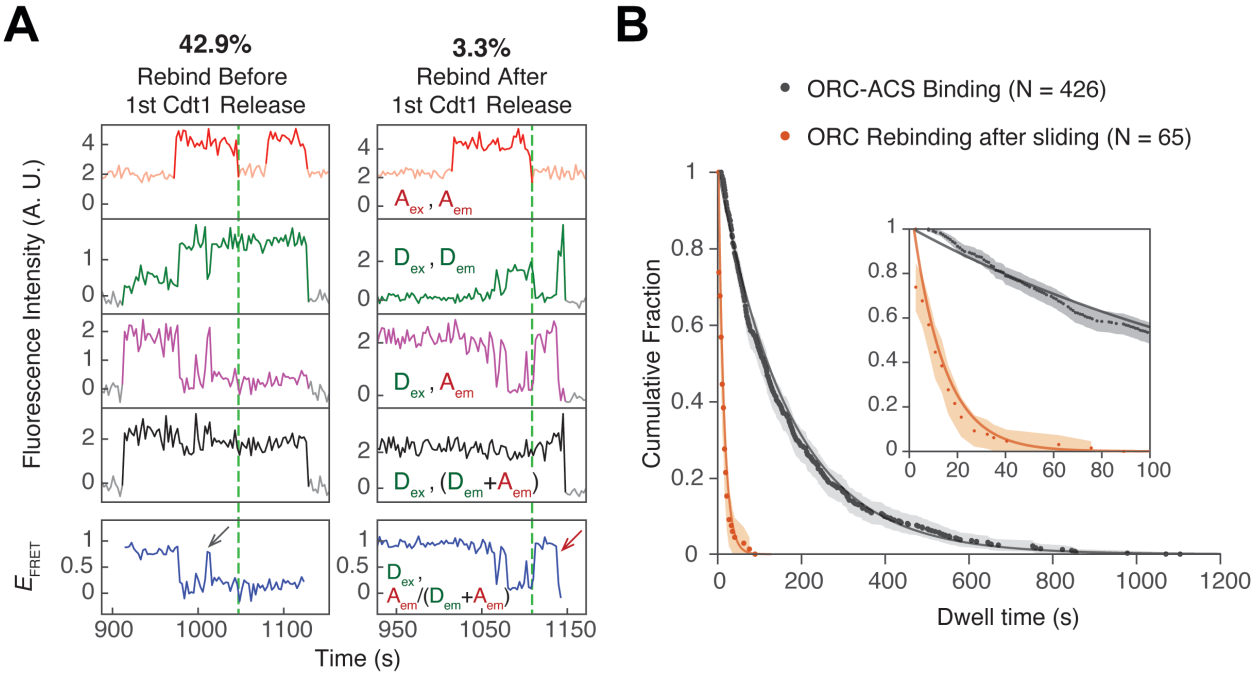


**Figure S11. ORC frequently and briefly rebinds the ACS after sliding begins.**

(A) Representative records from the ORC$^{N}$•+-4 + Cdt1$^{N\text{-}649}$ experiment (same as Fig. 6E) monitoring ORC association with the ACS in the presence of labeled Cdt1. Among the 91 ORC molecules that recruited a Cdt1 molecule, 39/91 (42.9%) showed rebinding (grey arrow) after ORC sliding begins. Only 3/91 (3.3%) rebound ACS after Cdt1 release (red arrow).

(B) ORC-ACS rebinding events are short-lived. The dwell time of ORC-ACS rebinding after ORC sliding (orange with 95% CI shading) is significantly shorter than ORC dwell time on ACS in the absence of other helicase-loading proteins (black with 95% CI shading). The ORC-ACS binding data (black) are from the ORC$^{N}$•-4 experiment (shown in *SI Appendix*, Fig. S2), in which the dwell times of the high $E_{\text{FRET}}$ state (indicating ORC-ACS interaction) are plotted. Inset: zoomed-in view of the dwell time survival plot. Solid curves: single-exponential fits (*SI Appendix*, Table S1).

### Zhang et al., Supplementary Tables

| **Apparent dissociation rate: Single exponential**<br>$k * exp(-k * (t - t_{min}))$ | | | | |
|---|---|---|---|---|
| **Fig** | **Dissociation of** | ***k*** **(s$^{-1}$)** | $t_{min}$ | **N** |
| S1C-D | ORC$^{C}$-DNA | 0.0131 ± 0.0005 | 2 | 1,108 |
| S1C | ORC$^{C}$-DNA + Cdc6 | 0.0073 ± 0.0004 | 2 | 679 |
| S1E | ORC$^{C\text{-}BPR}$-DNA | 0.0849 ± 0.0155 | 2 | 1,305 |
| S11B | ORC$^{N}$-ACS | 0.0059 ± 0.0003 | 2 | 426 |
| S11B | ORC$^{N}$(post-sliding)-ACS | 0.0763 ± 0.0121 | 5.4 | 65 |

| **ORC$^{C}$•+51 $E_{FRET}$ distributions: Single-component Gaussian**<br>$\frac{1}{\sigma\sqrt{2\pi}}\exp\{-\frac{(E_{FRET} - \mu)^2}{2\sigma^2}\}$ | | | | | |
|---|---|---|---|---|---|
| **Fig.** | **Proteins** | **μ** | **σ** | **N** | $n_t$ |
| 1D | ORC$^{C}$ | 0.62 ± 0.001 | 0.10 ± 0.002 | 186 | 25,388 |
| S1C | ORC$^{C}$+Cdc6 | 0.63 ± 0.001 | 0.08 ± 0.002 | 261 | 42,158 |
| S1E | ORC$^{C\text{-}BPR}$ | 0.53 ± 0.002 | 0.12 ± 0.002 | 1,305 | 19,940 |

| **MCM•+51 $E_{FRET}$ distributions: Two-component Gaussian**<br>$\frac{1}{\sqrt{2\pi}}\{\frac{p_{low}}{\sigma_{low}}exp[-\frac{(E_{FRET} - \mu_{low})^2}{2\sigma_{low}{}^2}] + \frac{(1-p_{low})}{\sigma_{high}}exp[-\frac{(E_{FRET} - \mu_{high})^2}{2\sigma_{high}{}^2}]\}$ | | | | | | | | |
|---|---|---|---|---|---|---|---|---|
| **Fig** | **Time interval** | **N** | $n_t$ | $p_{low}$ | $\mu_{low}$ | $\mu_{high}$ | $\sigma_{low}$ | $\sigma_{high}$ |
| 3C | 1 – 2 s | 135 | 540 | 0.81 ± 0.02 | *0.12 ± 0.006 | *0.55 ± 0.007 | *0.14 ± 0.005 | |
| | 10 – 11 s | | 540 | 0.10 ± 0.02 | | | | |
| 4B | TW1 | 130 | 520 | 0.68 ± 0.06 | 0.15 ± 0.01 | 0.49 ± 0.03 | 0.11 ± 0.01 | 0.11 ± 0.02 |
| | TW2 | | 1430 | 0.36 ± 0.05 | 0.17 ± 0.02 | 0.48 ± 0.01 | 0.10 ± 0.01 | 0.10 ± 0.01 |
| | TW3 | | 4550 | 0.37 ± 0.02 | 0.13 ± 0.01 | 0.36 ± 0.01 | 0.11 ± 0.01 | 0.19 ± 0.02 |
| | TW4 | | 6630 | 0.70 ± 0.05 | 0.23 ± 0.01 | 0.33 ± 0.02 | 0.13 ± 0.01 | 0.23 ± 0.02 |
| S8B | 0 - 10 s | 46 | 184 | 0.42 ± 0.07 | 0.20 ± 0.01 | 0.50 ± 0.01 | 0.11 ± 0.02 | 0.12 ± 0.03 |
| S8E | 0 - 10 s | 58 | 232 | 0.61 ± 0.09 | 0.21 ± 0.03 | 0.50 ± 0.05 | 0.08 ± 0.02 | 0.27 ± 0.03 |

| **Fit parameters** | |
|---|---|
| $k$ | Apparent dissociation rate ($k_{off\bullet app}$) |
| $t$ | Dwell time |
| $t_{min}$ | Minimum measureable dwell time |
| N | Number of colocalization events |
| $n_t$ | Number of frames during all colocalization events |
| μ | Gaussian mean |
| Σ | Gaussian standard deviation |
| $\mu_{low}$ | Gaussian mean of low $E_{FRET}$ component |
| $\mu_{high}$ | Gaussian mean of high $E_{FRET}$ component |
| $\sigma_{low}$ | Gaussian standard deviation of low $E_{FRET}$ component |
| $\sigma_{high}$ | Gaussian standard deviation of high $E_{FRET}$ component |
| $p_{low}$ | Proportion of low $E_{FRET}$ component |

**Table S1: Summary of fit parameters.**

Tables summarizing all fit parameters in this paper. Fit functions include single exponential, single-component Gaussian, and two-component Gaussian functions. The standard error is reported alongside each value, which is derived by bootstrapping (5,000 iterations). Parameters marked with asterisks (*) were fit globally to the two data sets. For dissociation rate fits, the fit parameter $t_{min}$ was included because only dwell times longer than 2 frames were considered to eliminate noise.

| FRET expt | Donor | Acceptor | Monitors | Data in Fig | Schematic in Fig |
|---|---|---|---|---|---|
| $ORC^{C}$•+51 | $ORC^{C}$ | *ARS1*$^{+51\text{-}Cy5}$<br>*ARS1*$^{+51\text{-}BHQ2}$ | DNA unbending | 1, 2 | 1B, 2A |
| MCM•+51 | Mcm2-$7^{3N\text{-}550}$ | *ARS1*$^{+51\text{-}Cy5}$ | DNA deposition | 3, 4A-B, 5 | 3A, S6A |
| MCM•-4 | Mcm2-$7^{3N\text{-}550}$ | *ARS1*$^{-4\text{-}Cy5}$ | MCM sliding leftwards from ACS | 4C-F | S6B |
| MCM•+82 | Mcm2-$7^{3N\text{-}550}$ | *ARS*$^{-+82\text{-}Cy5}$ | MCM sliding rightwards from ACS | 4C-F | S6B |
| $ORC^{N}$•-4 | $ORC^{N}$ | *ARS1*$^{-4\text{-}Cy5}$ | ORC•ACS interaction | 6A-E | 6A |
| $ORC^{N}$•+51 | $ORC^{N}$ | *ARS1*$^{+51\text{-}Cy5}$ | ORC sliding rightwards | 6F | S6C |

**Table S2. Summary of all FRET experiments in this study.**

| Sequence (5' → 3') | Used as | Final Product |
| --- | --- | --- |
| /5Biosg/GATCGGTGCGGGCCTCTTCGC | Biotinylated forward primer | All *ARS1* products |
| GGAAAGCGGGCAGTGAGCGC | Reverse primer for internally labeled fragment | All non-end labeled *ARS1* products |
| /5Alex488N/GGAAAGCGGGCAGTGAGCGC | End-labeled reverse primer | *ARS1*$^{+51\text{-BHQ2}}$ |
| /5Phos/AGTATTGTTTGTGCACTTGCCTGC | Reverse primer for biotinylated fragment | *ARS1*$^{-4\text{-Cy5}}$ |
| /5Phos/TTTCGTCAAAAATGCTAAG | Reverse primer for biotinylated fragment | *ARS1*$^{+51\text{-Cy5}}$, *ARS1*$^{+82\text{-Cy5}}$, *ARS1*$^{+51\text{-BHQ2}}$ |
| /5Phos/CATCTTGTTATTTTAC/iCy5/AGATTTTATGTTTAGATC | Internally labeled forward primer | *ARS1*$^{-4\text{-Cy5}}$ |
| /5Phos/GCAAGT/iCy5/GCACAAACAATACTTAAATAAATACTACTC | Internally labeled forward primer | *ARS1*$^{+51\text{-Cy5}}$ |
| /5Phos/CTTAAATAAATACTACTCA/iCy5/GTAATAACCTATTTCTTAGC | Internally labeled forward primer | *ARS1*$^{+82\text{-Cy5}}$ |
| /5Phos/GCAAGT/iBHQ_2/GCACAAACAATACTTAAATAAATACTACTC | Internally labeled forward primer | *ARS1*$^{+51\text{-BHQ2}}$ |
| CTTGTTATTTTACAGATTTTCTCCATTCTTCTTTTATGCTTGCAAAACAAAAGGCCTGCA | Nonspecific DNA in single-molecule experiments | 60 bp annealed nonspecific DNA |
| TGCAGGCCTTTTGTTTTGCAAGCATAAAAGAAGAATGGAGAAAATCTGTAAAATAACAAG | | |

**Table S3. List of oligos used in this study.**
